# 3D ultrasound fascicle tractography for objective muscle architecture analysis

**DOI:** 10.64898/2026.08.31.746736

**Authors:** Paolo Tecchio, Lara Schlaffke, Bart Bolsterlee, Daniel Hahn, Brent J. Raiteri

## Abstract

Muscle architecture shapes muscle function and changes with age, growth, training and disease, yet quantifying three-dimensional (3D) muscle architecture in vivo remains challenging. We introduce a hybrid fascicle tractography approach for freehand 3D ultrasound data that accurately reconstructs 3D muscle fascicles with respect to an objective, anatomically relevant coordinate system defined by the muscle’s central aponeurosis. The hybrid approach combines Hessian-based fascicle detection with wavelet-based refinement to generate volumetric fascicle orientations. In a synthetic dataset with known ground truth, fascicle orientations and lengths were estimated with errors of ≤2° and ∼1.5%, respectively. *In vivo*, the approach detected physiologically plausible fascicle lengthening in the human tibialis anterior following a passive plantar flexion rotation, whereas diffusion tensor imaging of the same muscle did not. The proposed method enables anatomically relevant, objective and non-invasive quantification of 3D muscle architecture *in vivo*, providing a practical framework for applications in clinical and applied muscle physiology.

## 1. Introduction

For over five centuries, scientists and artists have sought to visualize the musculoskeletal system and the internal structure of muscles beneath the skin. Indeed, muscle architecture – the spatial arrangement of muscle fibers and their connective tissues (Gans, 1982) – has been studied since the anatomical illustrations of the Renaissance (Vesalius, 1543), and was later described biomechanically by Borelli in the 17^th^ century (Borelli et al., 1680). Muscle architecture is a key determinant of a muscle’s function (Gans, 1982) that sets muscle-based limits of locomotor performance and the cost of transport (Clemente et al., 2024). Furthermore, muscles are highly plastic across growth (Chow et al., 2025), aging (Narici et al., 2021), training (Blazevich, 2006; Seynnes et al., 2007) and disease (Wang et al., 2024), making muscles promising non-invasive biomarkers of a person’s overall health, which is likely relevant for health screening and longitudinal monitoring. Although skeletal muscles can be readily imaged non-invasively to quantify local and two-dimensional architectural features such as fascicle length and fascicle angle, directly measuring an individual muscle’s complete three-dimensional architecture remains challenging (Binder-Markey et al., 2023; Dick and Hug, 2023).

Since the early attempts to understand how muscle architectural features shape muscle function and vary across muscles in human cadavers (Wickiewicz et al., 1983), advances in imaging technologies have revolutionized our ability to study muscle architecture *in vivo* and improved our understanding of muscle-tendon function (Cronin and Lichtwark, 2013). The main *in vivo* imaging modalities for quantifying muscle fascicle and fiber lengths, which affect a muscle’s force potential and metabolic energy expenditure (Bohm et al., 2019), are Brightness-mode (B-mode) ultrasound (US) and diffusion tensor imaging (DTI). DTI is an magnetic resonance imaging (MRI) protocol that is sensitive to the direction and magnitude of diffusion of water molecules in tissues that was first applied to track muscle fibers in 2002 (Damon et al., 2002). DTI offers moderate spatial resolution (order of millimeters) and can estimate three-dimensional (3D) muscle-fiber trajectories throughout the entire muscle volume, enabling non-invasive estimation of 3D muscle architecture (Damon et al., 2002). The *estimation* of 3D muscle-fiber architecture relies on the assumption that water molecules diffuse preferentially along the muscle fibers. Thus, DTI infers fiber trajectories indirectly from anisotropic water diffusion rather than measuring them directly (Damon et al., 2002). Moreover, the application of DTI to skeletal muscle is limited by the requirement for costly and time consuming high–field MRI (≥3T) scans to obtain adequate signal quality, as well as by the demanding and technically complex post–processing needed to derive reliable diffusion metrics (Forsting et al., 2022; Lockard et al., 2024).

B-mode ultrasound was first applied to measure human muscle fascicle lengths in the late 1990s (Fukunaga et al., 1997; Narici et al., 1996), and offers a cost-effective and portable alternative with higher spatial and temporal resolution (Gill, 2008). Yet, conventional two-dimensional (2D) US is constrained by a limited field of view and is susceptible to image noise, including speckle noise and refraction artifacts (Gill, 2008). Moreover, US is highly operator-dependent, particularly with respect to the orientation of the transducer when imaging muscle fascicles in the longitudinal plane (Bolsterlee et al., 2016). Traditionally, clinicians and researchers image muscle fascicles by aligning the transducer along a plane *assumed* to correspond to the fascicle plane (Narici et al., 1996; Rutherford and Jones, 1992). In this single 2D plane, muscle architecture is subsequently quantified manually or using automated fascicle detection algorithms (Ritsche et al., 2024; Seynnes and Cronin, 2020; Van Der Zee et al., 2025). However, recent DTI studies have shown that muscle fibers in many human muscles follow trajectories with curvatures in multiple planes, making it improbable to image fascicles in their entirety in a single 2D imaging plane (Bolsterlee et al., 2015; Suskens et al., 2023; Takahashi et al., 2022). To overcome this limitation, freehand three-dimensional ultrasound (3DUS) has been used to reconstruct entire muscle volumes and to estimate muscle architecture at rest and during contraction from 2D reconstructed planes (Barber et al., 2009; Bénard et al., 2011; MacGillivray et al., 2009).

For estimating muscle architecture from 3DUS data, many scientists rely on a single 2D reconstructed plane, subjectively oriented within the 3D muscle volume to measure fascicle angles and lengths (Andrews et al., 2024; Bénard et al., 2011; Cenni et al., 2018; Pincheira et al., 2022; Raiteri et al., 2016; Wang et al., 2023). In contrast, Rana et al. acquired multiple 2D scans in varying “longitudinal planes” and applied a wavelet analysis to estimate 3D fascicle orientations and lengths within the human triceps surae (Rana et al., 2013). More recently, Sahrmann et al. (2024) proposed the use of a single US sweep in the transverse plane and employed local curvature from the Hessian matrix to extract 3D fascicle orientations within the human tibialis anterior muscle (TA). Both approaches were validated using wire phantoms of known orientation. However, an important simplification was made during these validations; the wire orientation was either determined relative to the lab coordinate system (Rana and Wakeling, 2011) or relative to another set of wires (Sahrmann et al., 2024). Therefore, the wire phantom validation inherently solved one of the key difficulties with 3D fascicle detection; the definition of an objective anatomically relevant coordinate system. Although this 3D-detection problem was somewhat addressed for the image data by converting the 2D fascicle orientations to a 3D muscle-based coordinate system, such a coordinate system has arbitrary muscle thickness and width planes because the orientations of the muscle’s aponeuroses are ignored. Consequently, the accuracy of the two previous approaches to estimate 3D fascicle orientations relative to the muscle’s thickness and width planes remains unknown, and the “validity” of the image realignment step remains unclear.

Here, we aimed to leverage the single-sweep, freehand 3DUS method and develop a hybrid approach that combines the work of Rana et al. and Sahrmann et al. (Rana and Wakeling, 2011; Sahrmann et al., 2024) to reconstruct 3D muscle fascicles within an objective anatomical coordinate system. This objective anatomical coordinate system was based on a principal-component-analysis (PCA) derived central aponeurosis coordinate system of the human TA to derive meaningful muscle thickness and width planes (Raiteri et al., 2016). To test and validate our 3D fascicle reconstruction algorithm, we generated a synthetic fascicle volume and compared our 3D reconstructed fascicles with ground truth fascicle lengths and orientations. We deemed fascicle length errors of ±5% as acceptable and a median orientation error of ±2° as acceptable in both muscle thickness (XZ) and width (XY) planes (Rana et al., 2013; Sahrmann et al., 2024). To verify the accuracy of our hybrid approach, we also determined TA muscle volumes and 3D reconstructed fascicle lengths at two ankle angles *in vivo* from the left and right legs at rest (N=5, 10 legs) and compared the outcomes with those derived from the current gold standard of DTI fiber tractography, applied to the same legs. Based on previous work (Herbert et al., 2002), we expected fascicle or fiber lengthening of 5.8 mm following a 20° passive rotation of ankle plantar flexion.

## 2. Methods

### 2.1 Participants

We collected data from both legs of five participants (two males, age: 33 ± 9.5 yrs, mean ± standard deviation; mass 69.4 ± 7.2 kg; height 1.73 ± 0.08 m; tibia length = 357 ± 20.3 mm). All participants gave written informed consent prior to participating in the study and reported no recent (<24 months) history of lower limb injuries or neuromuscular diseases. The experimental procedures were approved by the local Ethics Committee of the Faculty of Sport Science at Ruhr University Bochum (EKS V 2023_06.1). The study was conducted in accordance with the ethical principles outlined within the declaration of Helsinki with the exception that our study was not preregistered.

### 2.2 Synthetic fascicle volume

To validate the detection of fascicle orientation in the muscle thickness (XZ) and width (XY) planes and to test the accuracy of the 3D fascicle reconstructions, we generated a synthetic volume with a fully defined ground-truth geometry. The volume consisted of 300 transverse slices (408 x 512 pixels, thickness x width) and included a planar synthetic aponeurosis that served as the objective anatomical coordinate system. Five circular structures (radius 12 pixels), representing fascicles in the muscle width plane, were defined in each slice as foreground (white) pixels. Fascicles were modeled as foreground pixels against a null background to ensure a high signal-to-noise ratio for initial geometry validation. Across slices, the circle centers were incrementally translated along the longitudinal (X) and medio-lateral (Y) axes, forming tubular 3D fascicle structures (Sahrmann et al., 2024) with controlled orientations. This produced deterministic orientations of 64.8 degrees in the muscle thickness plane and 25.5 degrees in the muscle width plane, with a mean fascicle length of 575.4 mm. All spatial parameters were scaled using an isotropic factor of 0.25026 mm/pixel to match the spatial resolution of the resliced 3DUS data.

### 2.3 3D ultrasound acquisition

Freehand 3D ultrasound data were acquired using Stradwin (Treece et al., 2003) (v.6.0.2). Prior to acquisition, the temporal and spatial calibration between the ultrasound system (ArtUs EXT-1H, transducer LV8-5N60-A, B-mode, 8.0 MHz, Telemed, Vilnius, Lithuania, 60-66db) and the motion capture system was performed using a water-bath method (Prager et al., 1998). The motion capture system comprised of four cameras (Flex 3, 0.3 MPixels, latency 10ms, 3D accuracy ± 0.5mm, 100 fps, OptiTrack, NaturalPoint, Corvallis, OR, USA) positioned around and above the volume of interest (Figure 1). Each camera’s lens was adjusted manually to enhance marker detection. A custom 3D-printed frame with four 9 mm diameter reflective markers (stereolithography model available on <u>GitHub</u>) was attached to the US transducer and defined as a rigid body in OptiTrack. The US transducer had a field of view of 64 mm (0.1205 mm per pixel) and was calibrated for an image depth of 50 mm (root mean square error = 1.12 mm). Volumetric US data were acquired at 67.1 fps with the transducer oriented transversely relative to the longitudinal axis of the TA muscle, which extends from the fibular head to a distal insertion on the foot (Zielinska et al., 2021). Data collection was performed by two trained investigators (PT and BR), who translated the transducer along the muscle’s longitudinal axis at an approximately constant velocity with constant pressure while participants laid supine on a bench and were instructed to remain relaxed. Scans were performed at two specific ankle angles (15° and 35° plantar flexion), which were attained by altering the orientation of a 3D-printed foot adapter (<u>GitHub</u>) that both feet were secured to via two Velcro straps. Each scan took <30 s (see results 3.4), the ankle angle that was tested first was randomized, and scans at each angle were repeated when excessive jittering was present in the scan, and 1 min rest was interposed between scans. Abundant acoustic gel combined with bacteriostatic ultrasound gel pads (Parker Laboratories; 90 mm diameter × 20 mm thickness) was used to optimize acoustic transmission (Klucinec, 1996) and minimize tissue compression. Each scan spanned from the fibular head to approximately the ankle joint, ensuring coverage of the entire TA muscle belly.

**Figure 1.**
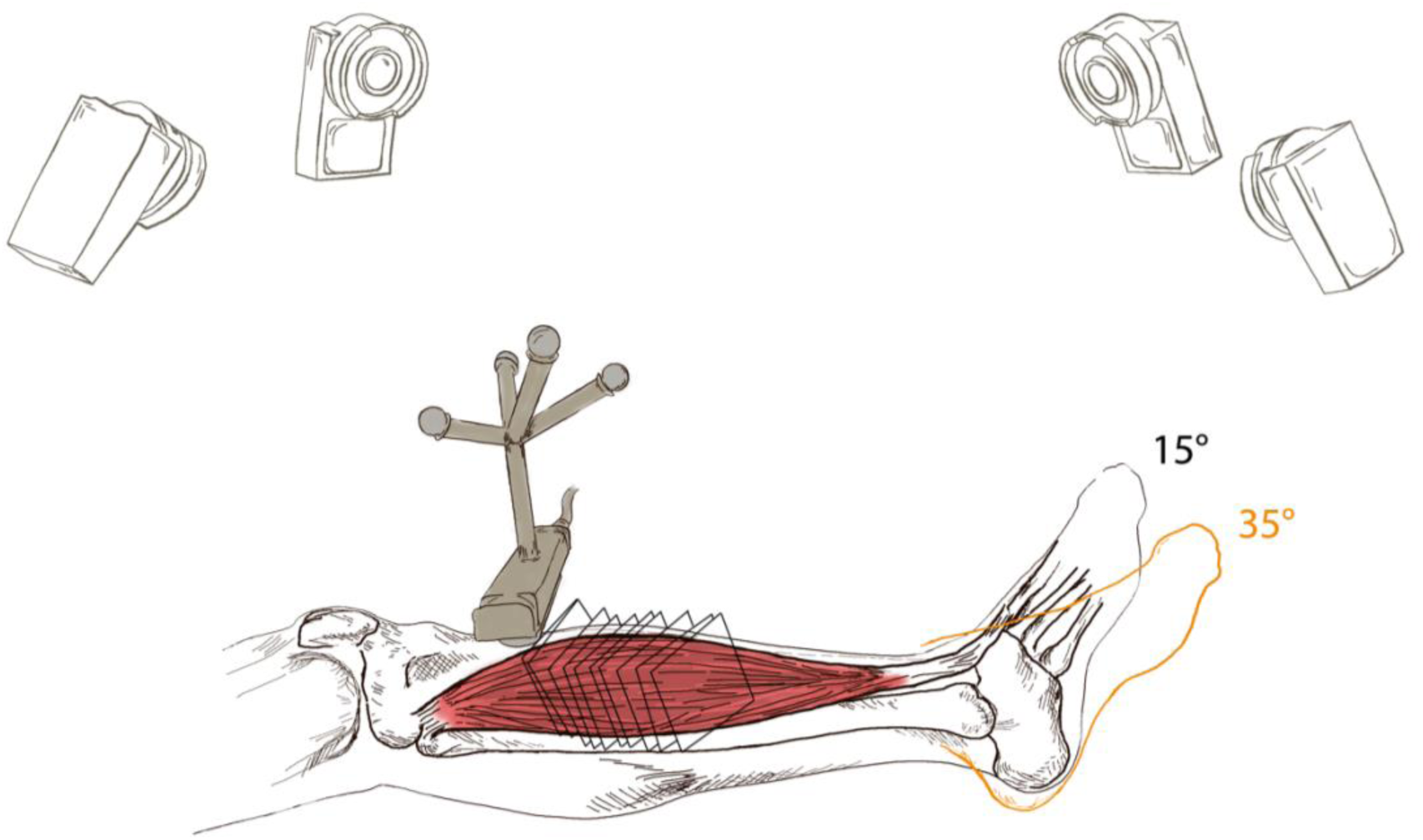
Schematic of the setup used to acquire freehand 3DUS data at two ankle angles (+15° and +35° PF, with 0° defined as 90° angle between the tibia and the foot sole).

### 2.4 3D ultrasound segmentation

All segmentations of the TA muscle and its central aponeurosis were performed by a single experienced investigator (PT; >200 h experience in US image segmentation). The TA was segmented by manually delineating the epimysial muscle borders from its proximal origin near the fibular head to approximately 1 cm distal to the termination of the central aponeurosis, where the TA’s deep compartment merges with the so-called “free” tendon. The borders of the central aponeurosis were also manually delineated throughout the volume, following the same procedure described in Raiteri et al. (Raiteri et al., 2016). Segmentation was performed using Stradview (v7.36). Surface reconstruction was performed using the regularized marching tetrahedra (Treece et al., 1999) with the software’s “high” resolution and “very high” smoothing settings enabled, and with hole-filling and open-surface correction options ticked. Following reconstruction, the investigator visually inspected the resulting surfaces throughout the entire muscle volume and, when necessary, refined the segmentation by adding or editing contours. The scattered 3D ultrasound data were subsequently resliced at an orientation that minimized the size of the output voxel data. The data was then exported as a constant grid (i.e., resliced) using a “two pixels” scaling factor, which provided a balance between spatial resolution and file size (isotropic voxel size: 0.25026 mm; file size ∼160 MB per file).

### 2.5 3D ultrasound fascicle reconstruction

The resliced image volume was then imported into MATLAB (The MathWorks Inc., MATLAB R2023b, Natick, Massachusetts, US) for 3DUS fascicle reconstruction. To reduce boundary-related artifacts that could impede accurate line detection, a dual-masking procedure was implemented (Sahrmann et al., 2024). For each transverse slice, the vertices defining the segmented TA contour were eroded toward its geometric centroid by 15% of the radial distance. Conversely, the central aponeurosis contour was dilated away from its centroid by 25% of the radial distance. Pixels outside the eroded TA boundary and inside the dilated aponeurosis boundary were masked (i.e. set to zero intensity; Figure 2a). 15% and 25% were chosen following visual inspection to safely remove any potential segmentation artifacts.

**Figure 2.**
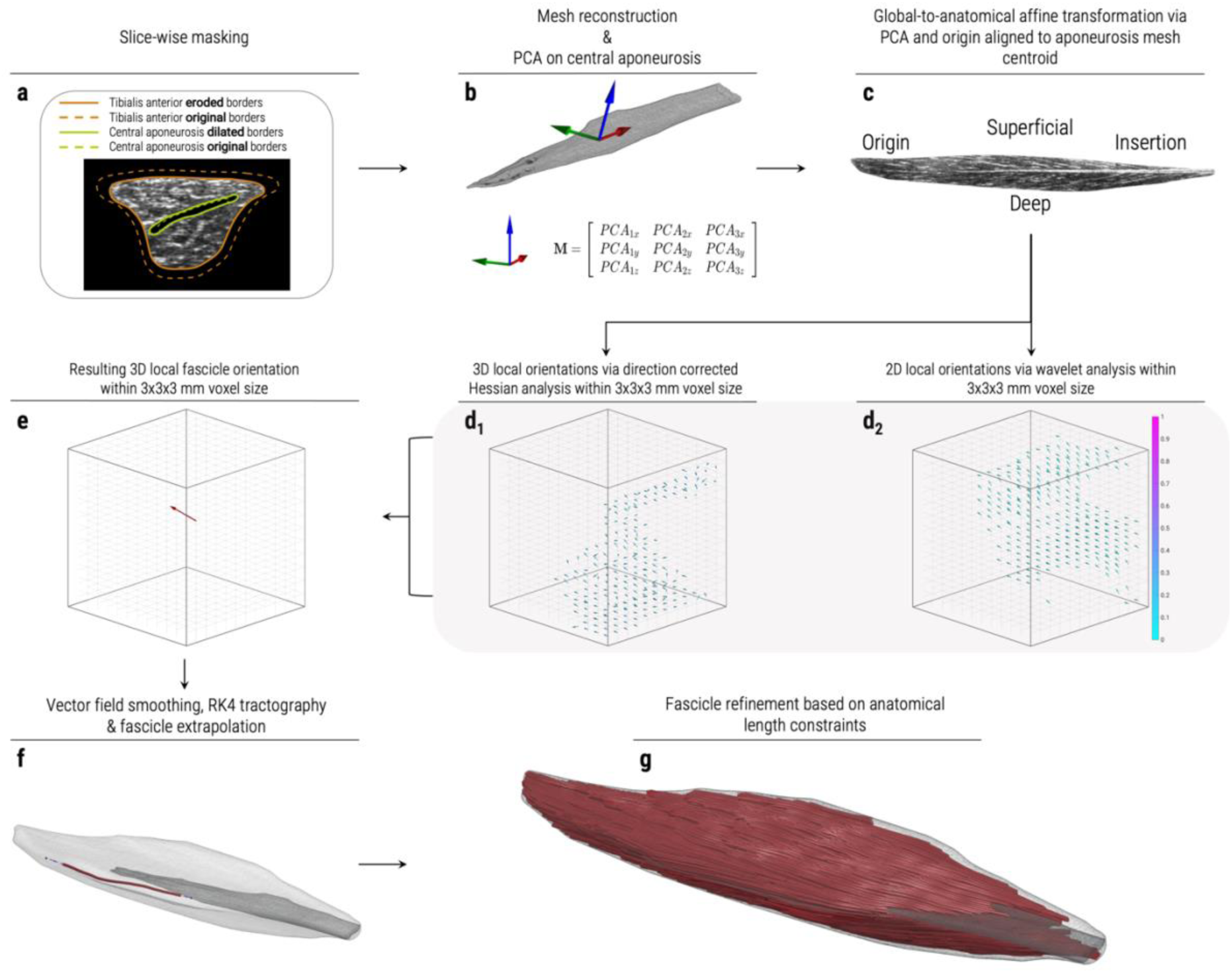
Pipeline workflow of the hybrid algorithm for 3D fascicle reconstruction. Zoom in for details.

The masked TA muscle volume was further partitioned into superficial and deep compartments. This compartmentalization was performed slice by slice by fitting a line of best fit (first-order polynomial) to the vertices defining the central aponeurosis within each transverse slice. The fitted line was used as a dividing boundary to separate voxels belonging to TA’s superficial and deep compartments (Hooijmans et al., 2025; Lansdown et al., 2007; Lockard et al., 2024; Sahrmann et al., 2024) . In the proximal region of the TA where the central aponeurosis was absent, the compartmental division was performed using the linear regression model obtained from the most proximal slice where aponeurosis vertices were available. Subsequently, the vertex coordinates of TA’s muscle belly and its central aponeurosis were used to reconstruct surface meshes via a crust algorithm (Giaccari, 2009). The resulting meshes were then processed using the *iso2mesh* package (Tran et al., 2020) to ensure watertightness (Figure 2b).

A weighted principal component analysis (PCA) was subsequently performed on the vertices of the central aponeurosis mesh (Raiteri et al., 2016). Because aponeurosis geometry varies primarily in length and width, the first, second and third PCA eigenvectors corresponded consistently to the longitudinal (X), medio-lateral (Y), and vertical (Z) axes, respectively (Figure 2b). As the sign of the PCA eigenvectors is inherently ambiguous, eigenvector directions were resolved by applying sign flips, when necessary, to enforce a consistent objective anatomical reference system. Specifically, eigenvectors were sign-flipped such that fascicles were visually oriented from positive to negative values along the longitudinal (X) axis (i.e. extending from right to left in the reconstructed longitudinal slices, from tendon insertion to muscle origin, see Figure 2c). This convention allowed fascicle angles to be expressed within the range of 90° to 270°, corresponding to the second and third Cartesian quadrants. The resulting sign-corrected eigenvectors were assembled into a 3 x 3 transformation matrix and applied to the TA and aponeurosis meshes, as well as to the image volumes. For the volumetric image data, affine transformations were implemented using the MATLAB function *imwarp* to account for shearing and reconstruction effects (Szeliski, 2022). Finally, both meshes and image volumes were translated to the centroid of the aponeurosis mesh (Figure 2b and 2c), which was defined as the origin of the objective anatomical coordinate system.

From here on, *U*, *V*, and *W* denote the 3D vector components of local fascicles within the PCA-defined anatomical reference system, corresponding to the longitudinal (X), medio-lateral (Y), and vertical (Z) axes, respectively. “*Orientations*” refer to the angles of the *U*, *V*, and *W* vectors in a specific plane (e.g., *θ* is the angle in the XZ plane), while the components describe the projections of each vector along each Cartesian axis.

Once the meshes and image volumes were within the objective anatomical coordinate system defined by the central aponeurosis of the TA, local 3D fascicle components were determined using the Hessian-based method proposed by Sahrmann et al. (Sahrmann et al., 2024), which is designed to identify tubular structures within image volume data (Figure 2d_1_). Components estimation was performed independently on each image volume corresponding to the superficial and deep compartments of the TA. To ensure consistency with the objective anatomical reference system defined by the PCA, the sign of each component was standardized such that its longitudinal (X) component was negative, corresponding to a distal-to-proximal fascicle direction.

To improve fascicle detection in noisy data and refine the *U* and *W* components, we additionally estimated local fascicle orientations in the muscle thickness (XZ) plane (i.e. fascicle angles) using a published wavelet-based fascicle detection algorithm (Kilpatrick et al., 2023). This 2D approach is particularly robust for detecting tubular structures in noisy images and provides supplementary information to ensure correct orientation sign assignment, which is finalized in a subsequent step. The algorithm was adapted for fully automated processing, with wavelet parameters optimized according to prior recommendations (Kilpatrick et al., 2023; Rana et al., 2013, 2009) and preliminary evaluations using the Hough transform. The adapted algorithm was applied to each slice oriented according to the PCA-based anatomical reference system to compute local orientation (*U* and *W* components) of fascicles (Figure 2d_2_). Processing was performed separately on TA’s superficial and deep compartments.

A user-defined voxel grid with a size of 3 × 3 × 3 mm^3^ was generated to subdivide each image volume. Each 3 mm voxel contained approximately 1723 of the original 0.25026 mm isotropic voxels. This voxel size was selected to ensure that a sufficiently large number of original voxels contributed to each local estimate, providing robust and numerically stable orientation calculations in noisy ultrasound data, while simultaneously minimizing redundancy and avoiding excessive spatial smoothing, particularly along the medio-lateral (Y) direction. The voxelization was applied across the entire image volume using overlapping blocks with a step size equal to 50% of the user-defined voxel size (i.e. 1.5 mm in this case). Each original voxel contained an orientation (i.e. fascicle angle in the muscle thickness [XZ] plane) and a convolution value from the wavelet analysis, or Nan (i.e. not a number) if no fascicle was detected. Within each 3 mm voxel, local fascicle components *U* and *W* were computed as a weighted mean of the original voxel size orientations. The weight *w_c_* for each voxel was derived from the wavelet convolution values, similar to Rana et al. (2013).

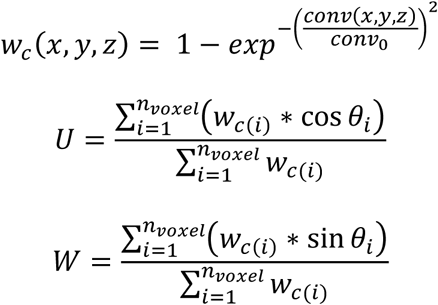

where *conv(x,y,z)* is the convolution value at a particular voxel; *conv_0_* represents the maximal convolution value over the entire image volume; {*x, y, z*} determine the 3D location of the user-defined voxel; *_θ_* represents the local fascicle angle detected by the wavelet analysis; *U* and *W* represent the weighted averaged local fascicle components along the longitudinal (X) and vertical (Z) axes, respectively.

Local 3D fascicle vector components (*U*, *V*, *W*) from the Hessian-based method were first computed as the median of all components within each previously user-defined voxel. The median was chosen over the mean because it provides a more robust estimate in noisy ultrasound data, reducing the influence of outliers. Before merging the Hessian- and wavelet-derived components within the user-defined voxel, the orientation *ϕ* representing the lateral component *V* from the Hessian-based method was calculated using the arctangent of the *V* and *U* components (i.e. angle in the muscle width [XY] plane). The *U* and *W* components from both methods were then combined using a weighting factor α ∈ [0,1]:

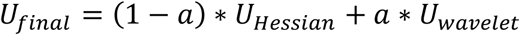

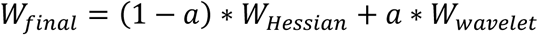

A high α value indicates increased confidence in the wavelet-derived components. For the entire analysis we used an α of 0.95. The α parameter was empirically chosen via direct observation and prior experimental experience. Following this combination, the lateral component *V* was reconstituted as:

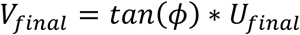

The resulting 3D vector components (*U_final_, V_final_, W_final_*) representing the local 3D fascicle vector in the user-defined voxel was normalized to unit length by dividing each component by the Euclidean norm (Figure 2e).

Following the calculation of the vector field within the user-defined voxel grid, the vector fields from the superficial and deep compartments were first combined. This was done by averaging overlapping components in regions where both compartments contribute vectors, particularly in the proximal part of the muscle, to create a single, unique 3D vector field. Vectors intersecting the aponeurosis or located within 1.5 mm of the aponeurosis mesh (i.e. half a voxel width) were then discarded. The resulting vector field was refined using a fast, unsupervised, and robust discretized spline smoother, as described by Sahrmann et al. (Sahrmann et al., 2024). This method performs a penalized least-squares fit to smooth the vector field. A conservative smoothing parameter of 0.1 was used, as averaging the components within the user-defined voxels had already reduced variability and noise.

### 2.6 Fascicle tracking

A normalized vector magnitude map (*M*) was computed based on the smoothed vector field as:

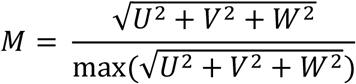

The normalization term max was taken over the entire vector field volume. This metric does not correspond to fractional anisotropy itself as defined in DTI, but instead provides a normalized estimate of local orientation consistency derived from the reconstructed vector field.

All voxels containing valid vectors were then used as seed points for bidirectional 3D fascicle tractography, performed using a 4^th^-order Runge-Kutta integration method (Butcher, 1996). Tracking parameters were as follows: step size = 1 mm, maximum number of steps = 150, *M* threshold = 0.1, and maximum turning angle = 40° (defined within a range of 0–90°). Fascicle endpoints were then either linearly extrapolated or truncated to the nearest intersection within the TA muscle and central aponeurosis meshes, with extrapolation limited to a maximum of 15 mm at each end (Bolsterlee et al., 2015) (Figure 2f). Fascicle length was then computed as the cumulative three-dimensional Euclidean distance along each reconstructed fiber. Only fascicles with lengths between 20 mm and 200 mm were retained (Figure 2g and Figure 3).

**Figure 3.**
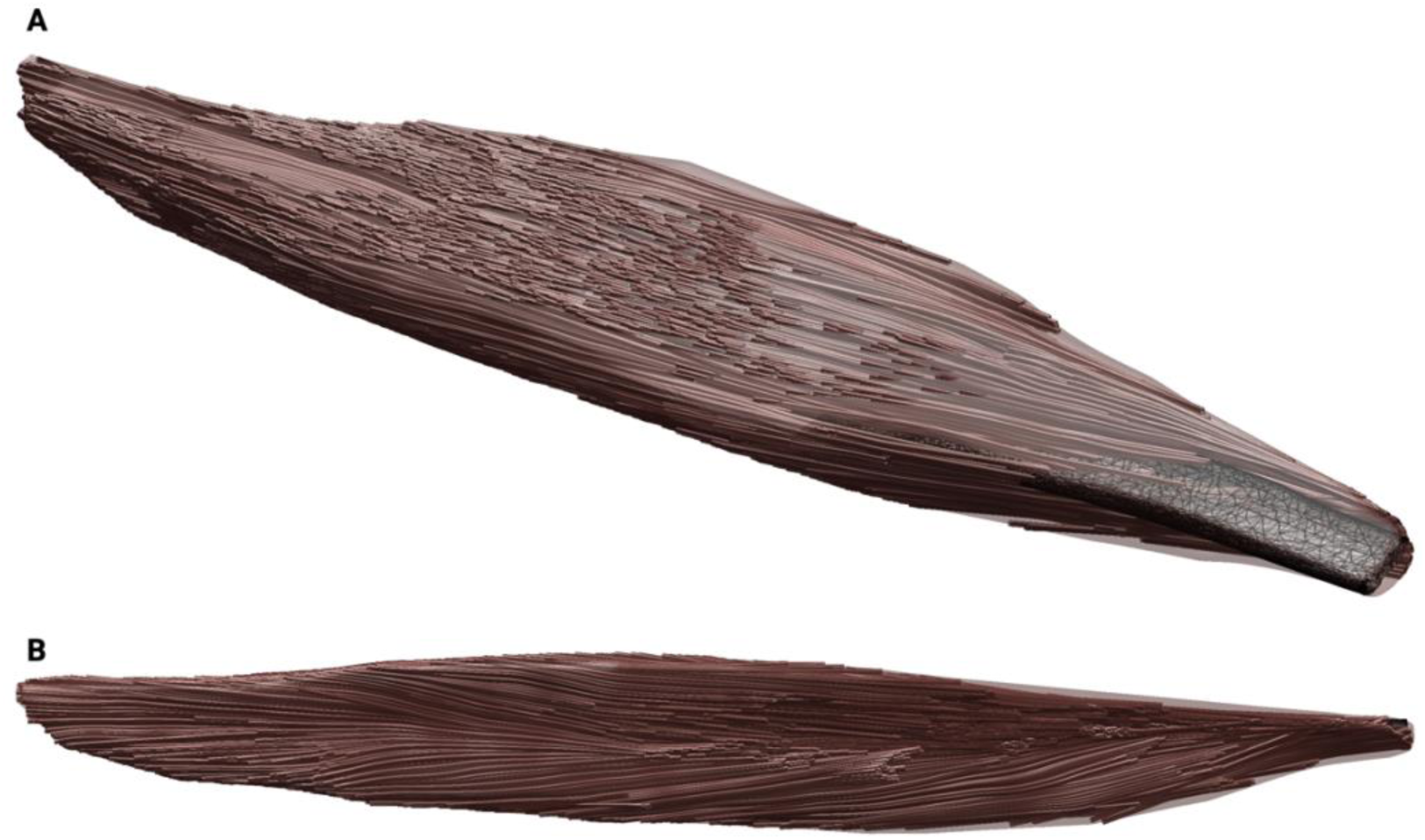
Example of reconstructed three-dimensional muscle fascicle streamlines from 3D ultrasound data acquired at 15° plantar flexion in one participant. The top panel (A) shows a three-dimensional view of the fascicle trajectories, while the bottom panel (B) shows a projection onto the sagittal plane (XZ) defined by the PCA-oriented aponeurosis coordinate system.

### 2.7 MRI/DTI

Magnetic resonance imaging (MRI) was performed on a 3T scanner (Prisma 3T, Siemens Medical Systems). Participants were positioned supine, and a Siemens Spine 32 Direct Connect coil in combination with a 18 channel body coil which was placed around the lower legs. Each scan lasted approximately 10 min. Three stacks of 15 cm in z-direction with an overlap of 3 cm were acquired to image the complete TA muscle belly, resulting in ∼30 min of total scanning per ankle joint angle. The protocol included Dixon, T2 mapping, and diffusion-weighted imaging sequences to characterize muscle structure. Dixon imaging was used to quantify intramuscular fat and water content, while T2 mapping provided information on water content and potential inflammatory processes and edema. A diffusion-weighted spin-echo echo-planar sequence with 43 gradient orientations at 6 b-values (b_0_=11, b_50_ = 3, b_100_ = 3, b_200_=6, b_400_ = 8, b_600_=12) was acquired to assess water mobility, and diffusion tensor imaging (DTI) was specifically performed at the same two ankle angles (15° and 35° plantar flexion) as for the 3DUS scans, using the same 3D-printed foot adapter. These DTI data were used to compare muscle volumes, fascicle length distributions, and fascicle orientations among joint positions and between imaging modalities. The voxel size was 3×3×6 mm³.

After the three image stacks obtained at each angle, images were merged in a single file, automated muscle segmentation with subsequent manual refinement and fiber tractography were performed using QMRITools (Froeling, 2019). Fiber tracts were not constrained to end on the central aponeurosis of the TA. The parameters used to perform tractography were an angle threshold of 30°, FA between 0.05 to 0.65, and a mean diffusivity between 0.5 to 2.5 x 10^-3^ mm^2^/sec and fiber lengths of 10 to 200 mm (defaults within QMRITools).

### 2.8 Statistics

For the synthetic fascicle volume, fascicle orientations in the muscle thickness (XZ) and width (XY) planes and fascicle lengths were compared with the ground truth using the median as the central tendency and the 25^th^–75^th^ percentile range to describe variability. For the *in vivo* data, two-way repeated-measures ANOVAs were performed with joint angle (15° and 35°) and imaging method (3DUS vs. DTI) as within-subject factors. Median fascicle length from the 3DUS reconstructions per participant was used in the analysis, whereas the mean fiber length from the DTI reconstructions was used as this is the output of QMRItools. Results are reported as the between-subject mean ± standard deviation for each ankle angle. The two-tailed significance level was set at 5%.

## 3. Results

### 3.1 Reconstruction and validation of ground truth synthetic data

We first validated the accuracy of our 3D reconstruction approach in detecting known fascicle orientations and lengths. To achieve this, we applied our hybrid approach, which combines two existing (Hessian- and wavelet-based) approaches, to a fully deterministic synthetic muscle volume containing five tubular fascicles embedded within a planar aponeurosis reference system. The imposed fascicle orientation was 64.8 degrees in the muscle thickness (XZ) plane and 25.5 degrees in the muscle width (XY) plane, with a mean fascicle length of 575.4 mm. The 3D reconstructed fascicle path was computed using 4^th^-order Runge-Kutta integration to obtain Euclidean length. This controlled scenario allowed quantification of 3D fascicle orientation and length reconstruction errors independent of biological variability.

Using a user-specified isotropic voxel size of 2 mm, our approach yielded a median relative error of −2.2 degrees [25^th^-75^th^ percentiles: −3.5 to −1.3°] in the muscle thickness plane and −0.3° [-0.9 to 1.5°] in the muscle width plane. Increasing the isotropic voxel size to 3 mm reduced the relative error further to −1.7° [−2.5 to −1.1°] in the muscle thickness plane, while remaining accurate in the muscle width plane (0.5° [−0.2 to −1.3°], Figure 4). Notably, the larger voxel size narrowed the distribution of fascicle orientations, indicating improved precision with spatial averaging. Fascicle length reconstructions were also accurate using 3 mm voxel size, as our approach reconstructed lengths of 566.6 mm [557.3 to 573.9 mm] from 193 successfully reconstructed fascicles, with an absolute error of 8.8 mm or ∼1.5% relative to the ground truth of 575.4 mm. Therefore, our hybrid approach achieved fascicle orientation errors within ±2° and length errors of approximately 1.5%, which is acceptable based on our a-priori criteria and verifies the validity of this hybrid approach under artificially-controlled conditions.

**Figure 4.**
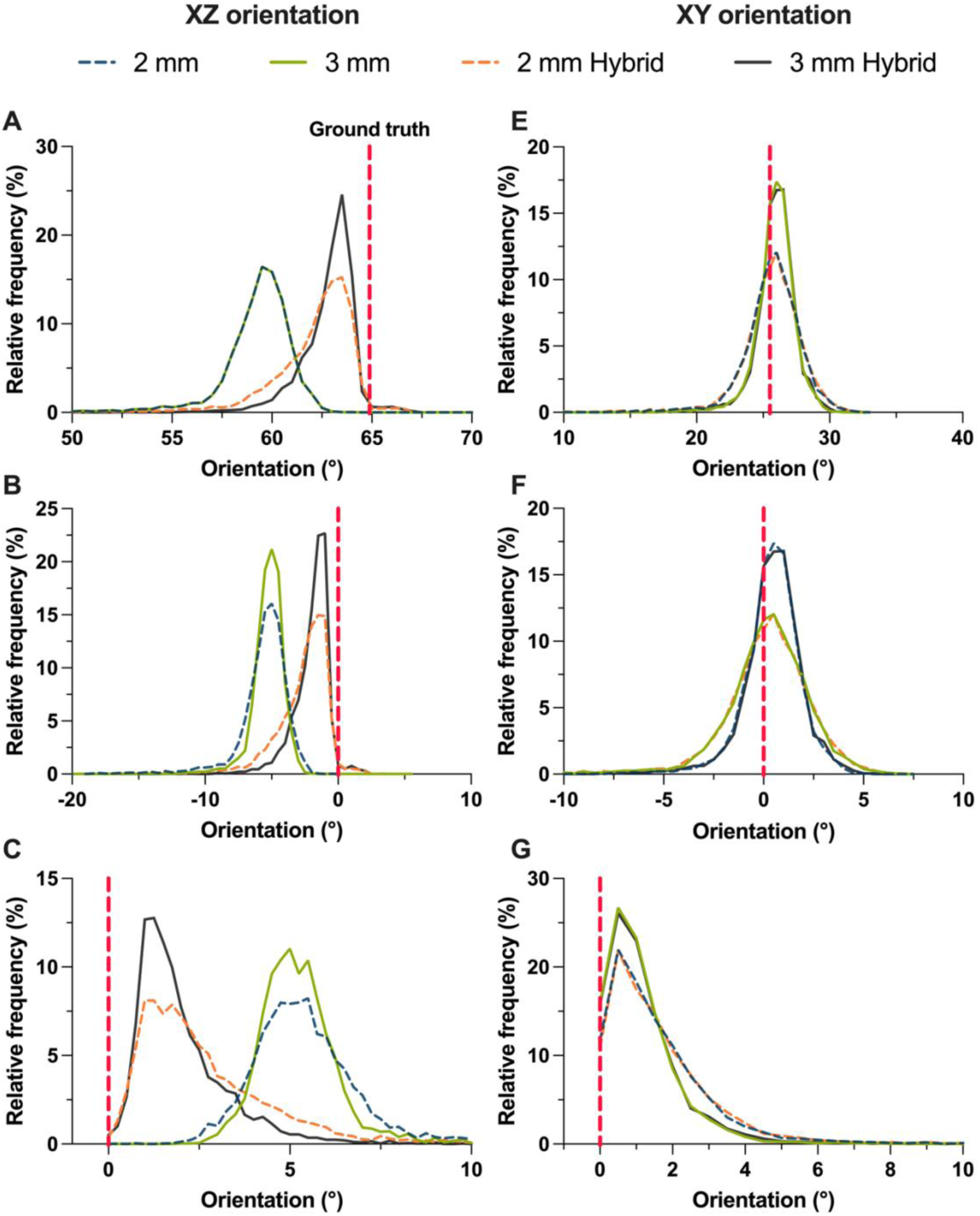
3D reconstructed fascicle orientation distributions in the muscle thickness (left) and width (right) planes calculated from the resulting vector fields. The 2 mm (blue) and 3 mm (green) traces show the median orientation distributions from the resulting vector fields using 2 and 3 mm voxel sizes and the Hessian analysis only (Sahrmann et al., 2024). The hybrid 2 mm (red) and 3 mm (black) traces show the median orientation distributions from the resulting vector fields using 2 and 3 mm voxel sizes and the combination of XZ vector field components from the wavelet analysis with the Hessian outcome. The first row (A & E) shows the actual fascicle orientations relative to the ground truth (vertical red dashed line), the second row (B & F) shows the relative fascicle orientation error compared with the ground truth, and the last row (C & G) shows the absolute maximum fascicle orientation error compared with the ground truth.

We further compared the performance of our hybrid approach with the standalone Hessian-based method (Sahrmann et al., 2024), under the same conditions and voxel sizes (Figure 4). Using 2 and 3 mm isotropic voxel sizes, the muscle thickness plane fascicle orientation errors were −5.3° [−6.2 to −4.5°] and −5.1° [−5.8 to −4.5°], respectively, while the muscle width plane fascicle orientation errors remained small (−0.5° [−1.3 to −0.3°] and 0.3° [−0.9 to 1.5°]). Although the Hessian-based method accurately estimated the fascicle orientations in the muscle width plane, this method alone exhibited a constant fascicle orientation bias in the muscle thickness plane, as previously reported (Sahrmann et al., 2024). Integrating wavelet-derived fascicle orientation information drastically reduced the fascicle orientation error in the muscle thickness plane, demonstrating superior spatial accuracy of our hybrid approach.

### 3.2 Influence of User-Defined Isotropic Voxel Size on Orientation Detection

Orientation detection from US images is affected by speckle noise, refraction artifacts and spatial heterogeneity. Therefore, we examined the effect of isotropic voxel size on the fascicle orientation detections. Small isotropic voxel sizes (e.g., 1 or 2 mm), although preserving the spatial variability, may lead to a relatively small number of native voxels (0.25026 mm isotropic size) within the analyzed block. For instance, the 1 mm isotropic voxel size has only 4*4*4 voxels or a total of 64 voxels per block. Conversely, large isotropic voxel sizes (e.g., ≥5 mm) may result in over-averaging within a spatially heterogenous block, causing a loss in spatial variability, which may lead to bias in the orientation detection. An isotropic voxel size of 3 × 3 × 3 mm provided a balance between reducing noise and preserving spatial heterogeneity (approximately 1,723 native voxels per block). Importantly, as the voxel size remains user-defined, on-the-fly adjustments are possible for different muscle architectures or different imaging resolutions. Tests on a real phantom dataset containing three curved wires in a water bath confirmed the trade-off between voxel size and spatial variability loss (Figure S12-15), where we found a 2 mm instead of a 3 mm isotropic voxel size better capture orientation heterogeneity in the width plane but results in redundant information. This highlights that voxel size selection requires careful consideration. Furthermore, in regions of steep changes in fascicle curvature (>∼20°), the Hessian-based orientation detection in the XY plane tends to underestimate local orientations (Figure S16).

### 3.3 *In vivo* performance and comparison with DTI

We applied our method to *in vivo* 3DUS data from the human TA muscle acquired at 15° and 35° of plantar flexion. Muscle volumes and fascicle lengths were compared with those obtained by DTI on the same legs. Over a 20° passive rotation of ankle plantar flexion, muscle volume was expected to remain unchanged, while fascicles/fibers should lengthen due to passive muscle-tendon unit elongation. Mean TA muscle volume derived from 3DUS was 105.7 ± 25.4 ml and 107.5 ± 26.9 ml at 15° and 35° of plantar flexion, respectively. Corresponding DTI-derived volumes were 122.6 ± 29.7 ml and 122.9 ± 30.3 ml, respectively (Figure 5A). Muscle volume was therefore unchanged over the range of joint rotation, but was different between the two techniques (*P* < 0.001). Median fascicle length derived from 3DUS increased from 93.6 ± 11.7 mm at 15° plantar flexion to 99.4 ± 10.5 mm at 35° plantar flexion (lengthening of 5.7 ± 3.3 mm, *P* < 0.001). Yet, DTI-derived mean fiber length decreased from 112.4 ± 17.0 mm to 106.1 ± 25.7 mm over the same passive plantar flexion rotation (shortening of 6.3 ± 11.5 mm, Figure 5B, *P* = 0.22). Thus, whilst 3DUS detected the expected fascicle lengthening with joint rotation (Herbert et al., 2002), DTI-derived tractography indicated fiber shortening. In addition, using participant-specific shank length and the previously reported value of 0.53 mm·deg^-1^ delta fascicle length and fascicle to tendon ratio change of 0.55 from Herbert et al.(Herbert et al., 2002), we predicted subject-specific fascicle length changes over 20° ankle plantar flexion. The mean predicted fascicle length change was 5.8 ± 0.3 mm.

**Figure 5.**
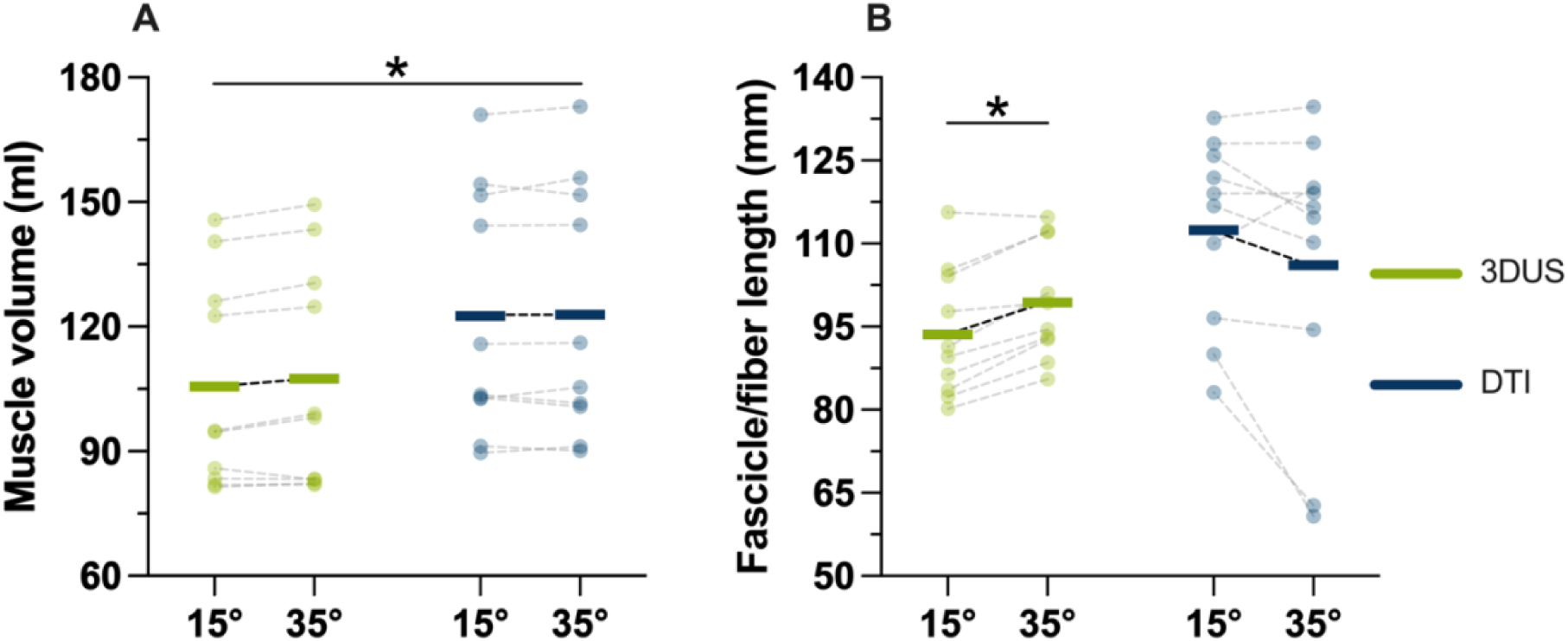
Individual tibialis anterior muscle volumes in milliliter (A) and muscle fascicle / fiber lengths (B) measured at two ankle angles (15° and 35° plantar flexion). Each dot represents one leg and measurements were obtained from both legs of five participants. Green dots and lines indicate outcomes estimated from 3D ultrasound (3DUS), while blue dots and lines indicate outcomes obtained from MRI/DTI data. The horizontal lines represent the means.

### 3.4 3DUS acquisitions details

The mean transducer sliding speed during 3DUS acquisitions was 10.1 ± 1.4 mm·s⁻¹ with a mean scan time of 24.5 ± 4.0 s, yielding >1300 frames per scan. Sampling density averaged 67 ± 9 frames·cm⁻¹ (Figure S5), exceeding the ∼33 frames·cm⁻¹ threshold needed for accurate voxel reconstruction and ensuring >80% detail recovery (Nikolaev et al., 2022). Transducer speed and frame rate were checked throughout data acquisition to maintain this threshold, a parameter often overlooked in previous freehand 3DUS studies.

## 4. Discussion

Accurate and reliable measurements of 3D muscle architecture *in vivo* have broad relevance. However, until this work, accurate and reliable *in vivo* quantification of 3D muscle architecture was performed relative to an arbitrary or user-defined plane (Dick and Hug, 2023), which limits the accuracy and usefulness of architecture outcomes as tools to track potential health changes (Chow et al., 2025; Moreau et al., 2010). The present hybrid approach provides a solution to this longstanding methodological barrier by allowing 3D muscle architectures to be objectively compared between people and over time, thus opening new avenues to use 3D architecture outcomes as reference data for monitoring muscle states and changes due to aging, growth, training or disease.

Our method outperforms previous methods as it combines Hessian-based fascicle detection with wavelet-based refinement and streamline tractography on 3D ultrasound data to generate accurate and reliable volumetric fascicle orientations and lengths. Another benefit of our method is that rather than relying on subjectively oriented or arbitrarily oriented 2D planes (Barber et al., 2009; Bénard et al., 2011; MacGillivray et al., 2009; Mantecón-Tagarro et al., 2025), 3D muscle architecture is reconstructed within an objective anatomical coordinate system with anatomically constrained muscle thickness and width planes. Consequently, the 3D muscle architecture outputs are anatomically relevant, as well as valid and biomechanically plausible, which will allow various future applications within clinical and applied muscle physiology.

We performed deterministic validation using a synthetic fascicle volume with known ground-truth orientations and lengths. Our hybrid approach reconstructed 3D fascicle lengths within ∼1.5% of the ground truth and 3D fascicle orientations within ±2° (median error) of the ground truth in both muscle thickness and width planes. Increasing user-defined isotropic voxel sizes from 2 to 3 mm reduced spatial noise in the fascicle orientation estimates without compromising accuracy in synthetic fascicle detection. Nevertheless, the isotropic voxel size parameter requires careful consideration and possible tuning in other conditions (e.g., active contractions) and in other muscles to ensure a balance between accuracy and robustness due to differences in fascicle orientation heterogeneity and ultrasound image noise.

Beyond our synthetic data validation, which had a favorable signal-to-noise ratio, we further stressed the hybrid approach using a physical phantom consisting of three curved wires submerged in a water bath. To challenge the algorithm, the wires were designed with steep slopes and non-physiological curvatures exceeding 30° in both frontal and sagittal planes (Figure S12-16). The algorithm successfully detected and partially tracked all three wires, though underestimation of local orientations in the frontal plane (XY) became apparent for local curvatures >20° (within3 mm isotropic voxel, Figure S16), while the sagittal (XZ) plane remained accurate. Although different filtering may improve local detection, excessive filtering can also introduce artifacts. Fortunately, such extreme local curvatures are rarely observed *in vivo* (Takahashi et al., 2022), and the abundance of parallel fascicles arrangement provides local redundancy that improves local orientation estimation.

*In vivo* application of our hybrid 3D fascicle reconstruction algorithm on the human TA revealed physiologically plausible muscle architectural outcomes. The 3DUS reconstructed fascicles lengthened by ∼5.7 mm as the ankle joint was passively plantar flexed from 15° to 35° plantar flexion. This ∼5.7 mm of fascicle lengthening closely aligns with theoretical predictions from Herbert et al. (Herbert et al., 2002), who reported muscle-tendon unit lengthening of 0.53 mm·deg^-1^ for the TA during passive plantar flexion rotations based on five cadaver studies. Based on this number, a 20° plantar flexion rotation results in 10.6 mm of TA muscle-tendon unit lengthening. As only 55% of this displacement is accommodated by TA’s muscle fascicles in a relaxed state based on *in vivo* 2D ultrasound data from six individuals (Herbert et al., 2002), the predicted fascicle length change is 5.83 mm, which closely agrees with our predicted fascicle length change of 5.79 mm based on our participant’s specific shank lengths. Although this estimate is based on 2D ultrasound data, fascicle length change estimates are more accurate than absolute length estimates (Day et al., 2017), and random errors in 2D transducer alignment should be mitigated when averaging length changes from a sample. This high level of agreement between the predicted fascicle length change and our 3DUS outcome suggests that our 3D fascicles reconstruction effectively captured the complex but expected interaction between contractile tissue and series elastic elements during TA muscle-tendon unit lengthening.

In contrast, the results from DTI tractography on the same participants indicated that TA muscle fibers shortened following the passive 20° ankle plantar flexion. The unexpected fiber shortening may be because of TA’s circumpennate architecture (Hirshowitz et al., 1987) with superficial and deep compartments along the distal two thirds of its belly length, which might introduce noise into DTI-based tractography when fiber tracts are not anatomically constrained to end on TA’s central aponeuroses and start at the superficial or deep muscle border (Bolsterlee et al., 2019). Notably though, even following work to improve stopping criteria in DTI tractography by considering tendinous tissues, test-retest reliability for fiber length measurements from the TA remained poor (Bolsterlee et al., 2019). Additionally, the repeatability of the fiber tract lengths from the TA remained the worst from four lower limb muscles with shortening sometimes observed when lengthening was expected (Oudeman et al., 2016), which indicates an inherent fiber tractography limitation when applied to the TA.

Aside from the fascicle and fiber length differences between 3DUS and DTI, there were discrepancies in muscle volume that likely arise from methodological differences. In particular, local transducer pressure during 3DUS acquisition may induce tissue deformation and therefore reduce muscle cross-sectional areas (Ryan et al., 2021), which lead to a lower volume compared with MRI-based segmentation. Furthermore, US images have less image contrast than MRI images, which can complicate the visual determination of muscle boundaries. Nevertheless, recent work (Ritsche et al., 2025; Sponbeck et al., 2021) found similar discrepancies between 3DUS and MRI derived TA muscle volumes within the same individuals, which could also be due to systematic differences in segmentation criteria.

Importantly, these volumetric differences were systematic at both ankle angles and did not affect our hybrid approach to detect biomechanically plausible differences in 3DUS fascicle lengths at different ankle angles.

The cost and accessibility of US imaging further enhances the potential and usefulness of our hybrid approach to estimate 3D muscle architecture. Compared with MRI-based methods, US can image with a higher spatial resolution at a faster speed, is non-claustrophobic, is easily accessible, and relatively inexpensive. Additionally, US can be used on patients with MRI-incompatible implants. The field of 3DUS acquisition itself is also constantly evolving, with emerging technologies such as deep learning reconstruction methods enabling volumetric imaging without the need for external trackers (Eid et al., 2025). Similarly, there has been substantial improvement with regard to artificial intelligence methods used for automatic segmentations (Rivera et al., 2025). The integration of these artificial-intelligence-based tools with the current hybrid approach proposed here, has the potential to provide even more automatic and faster reconstruction methods with minimal operator input. Such automation could enable complete volumetric muscle analysis directly at the doctor’s office, on the sports field, or at the bedside in hospital or at home. Therefore, a portable and completely automated 3D muscle architectural assessment tool is a realistic possibility for objective longitudinal monitoring of muscle architecture and health across diverse populations and environments.

### 4.1 Limitations

Some methodological limitations should be considered. The segmentation of the TA muscle and the central aponeurosis was done manually, although previous studies have indicated that this technique is highly reliable (Hanssen et al., 2023; Raiteri et al., 2016; Wang et al., 2023). The anatomical referencing system was based on the PCA of the central aponeurosis, but not all muscles possess a clearly defined internal aponeurosis. In such cases, the reference coordinate system could be based on the PCA axes derived from the muscle surface mesh itself (Rana et al., 2013; Sahrmann et al., 2024). However, because muscle cross-sectional areas vary along the muscle’s length and the muscle does not deform uniformly during passive (Herbert et al., 2019; Hodson-Tole et al., 2016; Moo et al., 2016) or active (Li et al., 2024; Pinto et al., 2023) (i.e. during contraction) conditions, the second and third PCA-derived axes will remain arbitrary and architecture will not be able to be accurately and reliably compared across conditions. Careful axis alignment and consistency checks would therefore be required when extending our hybrid approach to other muscles without a circumpennate architecture. Further, we performed only single-sweep 3DUS acquisitions. However, we expect comparable results with a sufficient spatial sampling density following multiple-sweep data acquisitions (i.e. >33 frames·cm⁻¹ of scan).

Importantly, our hybrid method was evaluated *in vivo* only under passive conditions at two joint angles with minimal passive tension. Yet, the robustness of the hybrid vector-field detection suggests that this approach is well suited investigating 3D muscle architecture at other joint angles and during isometric (i.e. fixed-end) muscle contractions. Such applications would enable the quantification of local and global muscle fascicle, belly and aponeurosis deformations *in vivo*. However, 3DUS currently remains constrained to static assessments. Lastly, to ensure consistency with the criteria used in the DTI QMRITool, we treated all voxels as potential seeds for tracking, rather than restricting the tracking to seeds adjacent to the central aponeurosis mesh. Although it is reasonable to assume that fascicles/fibers should originate at the central aponeurosis and terminate at the epimysium of the TA, we adopted this broader seeding strategy to enable a fair comparison by applying the same selection criteria as used in the 3DUS analysis.

## Data availability

The ultrasound and MRI/DTI data, scripts, statistics and extra material are available at the following GitHub repository: https://github.com/PaulT95/3DUS-Tractography-Study.

## Author contributions: CRediT

**P.T.:** Conceptualization, Investigation, Data curation, Formal analysis, Methodology, Visualization, Writing – original draft, Writing – review & editing.

**L.S.:** Investigation, Data curation, Formal analysis, Visualization, Writing – review & editing.

**B.B.:** Formal analysis, Writing – review & editing.

**D.H.:** Conceptualization, Supervision, Resources, Writing – review & editing.

**B.J.R.:** Conceptualization, Investigation, Data curation, Writing – review & editing.

## Funding

This work was partly supported by the German Research Foundation (DFG; 565338450).

## Competing interests

The authors declare no competing interests.

## Notes

https://github.com/PaulT95/3DUS-Tractography-Study

### Competing Interest Statement

The authors have declared no competing interest.

