## Supplemental material for "3D ultrasound fascicle tractography for objective muscle architecture analysis"

Synthetic fascicle volume

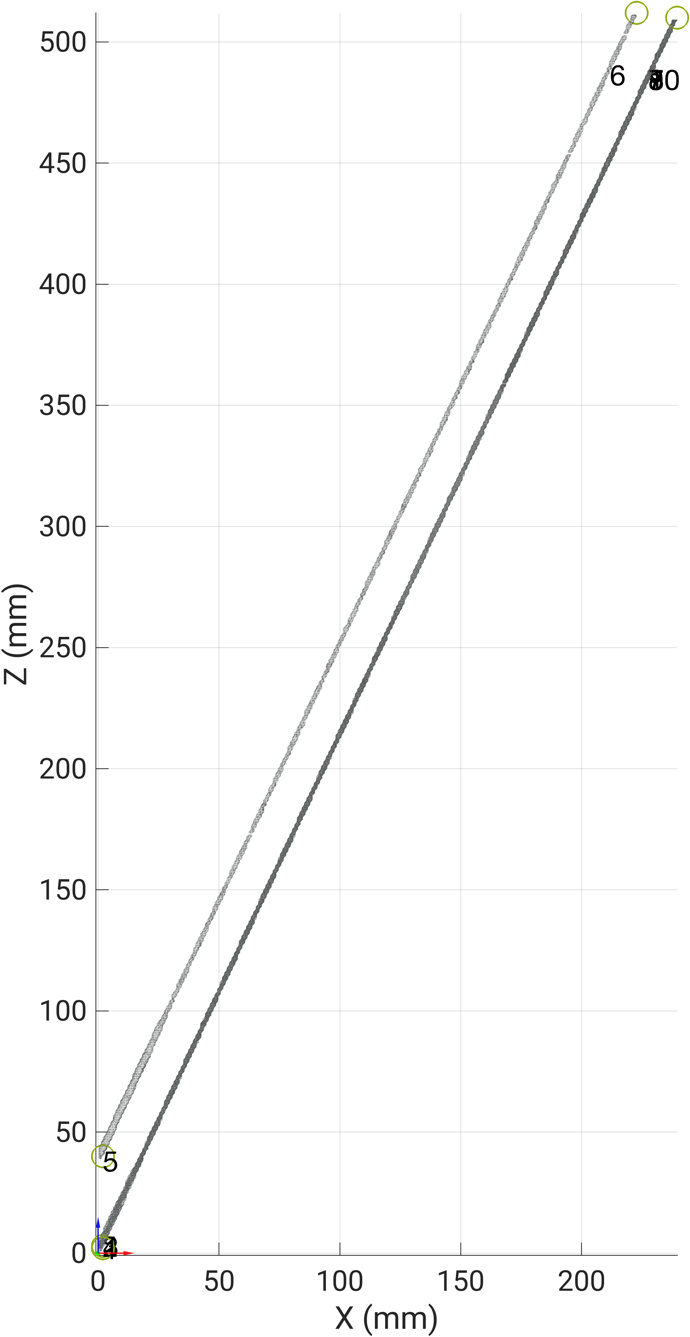

Figure S. 1. Parasagittal (XZ) plane projection of the synthetic fascicle volume based on the coordinate system of the aponeurosis. The volume has five tubular structures (each with a 12-pixel radius) modeled with a deterministic orientation of 64.8° in this XZ plane. This view highlights what would typically be captured by traditional 2D ultrasound methods that assume a single, fixed "fascicle plane" with correct parallel alignment of the ultrasound transducer’s longitudinal plane with the orientation of the central aponeurosis’ longitudinal plane. Presenting the fascicles with this view highlights consistent fascicle orientations in 2D, but this is not the case; fascicles actually move out of this plane due to curvature in the perpendicular frontal (XY) plane, necessitating the hybrid and anatomically-based 3DUS fascicle tracking approach developed in this study.

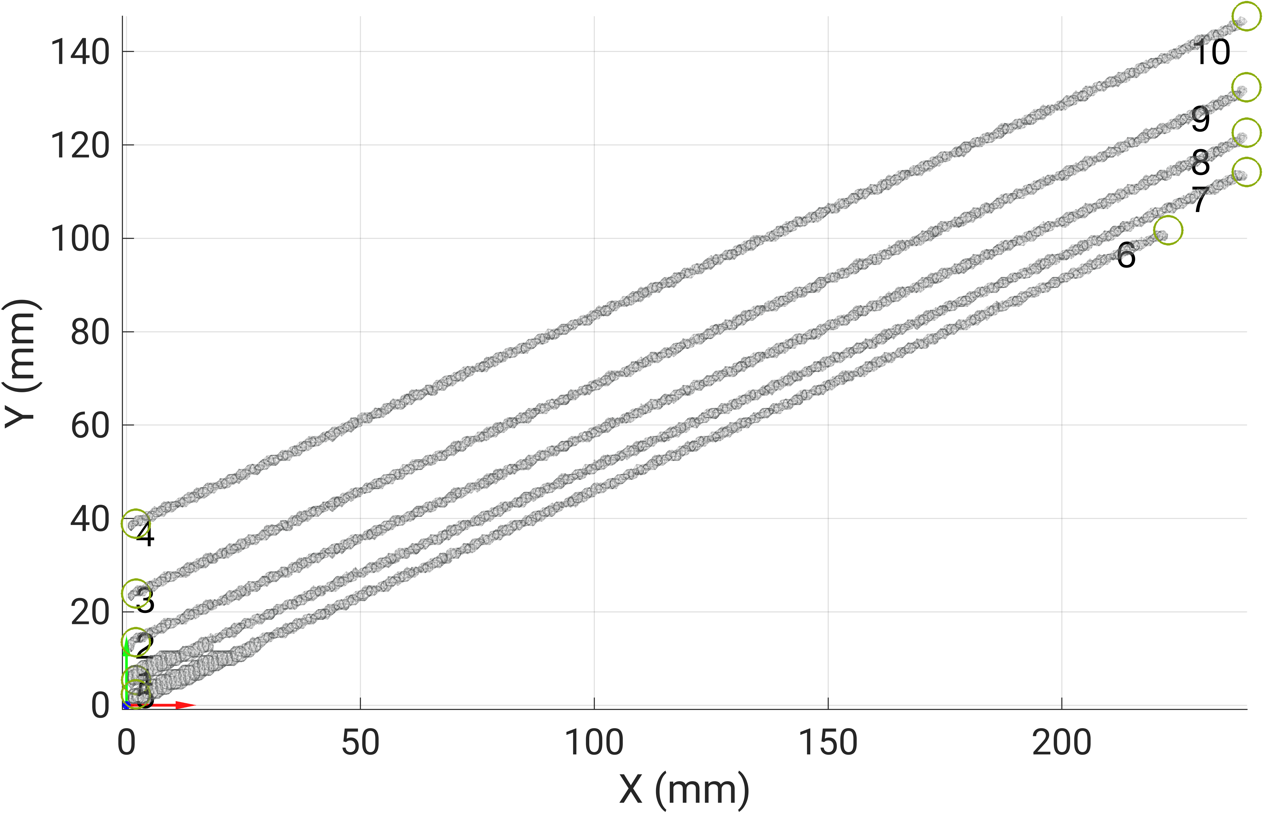

Figure S. 2. Frontal or medio-lateral (XY) plane projection of the synthetic fascicle volume based on the coordinate system of the aponeurosis. The visualization demonstrates how the modeled fascicles span the muscle width plane relative to the planar synthetic aponeurosis, providing a ground-truth reference for validating frontal plane curvature measurements (i.e. fascicle orientation detection in the muscle width plane). In this view, fascicles were modeled with a deterministic orientation of 25.5°.

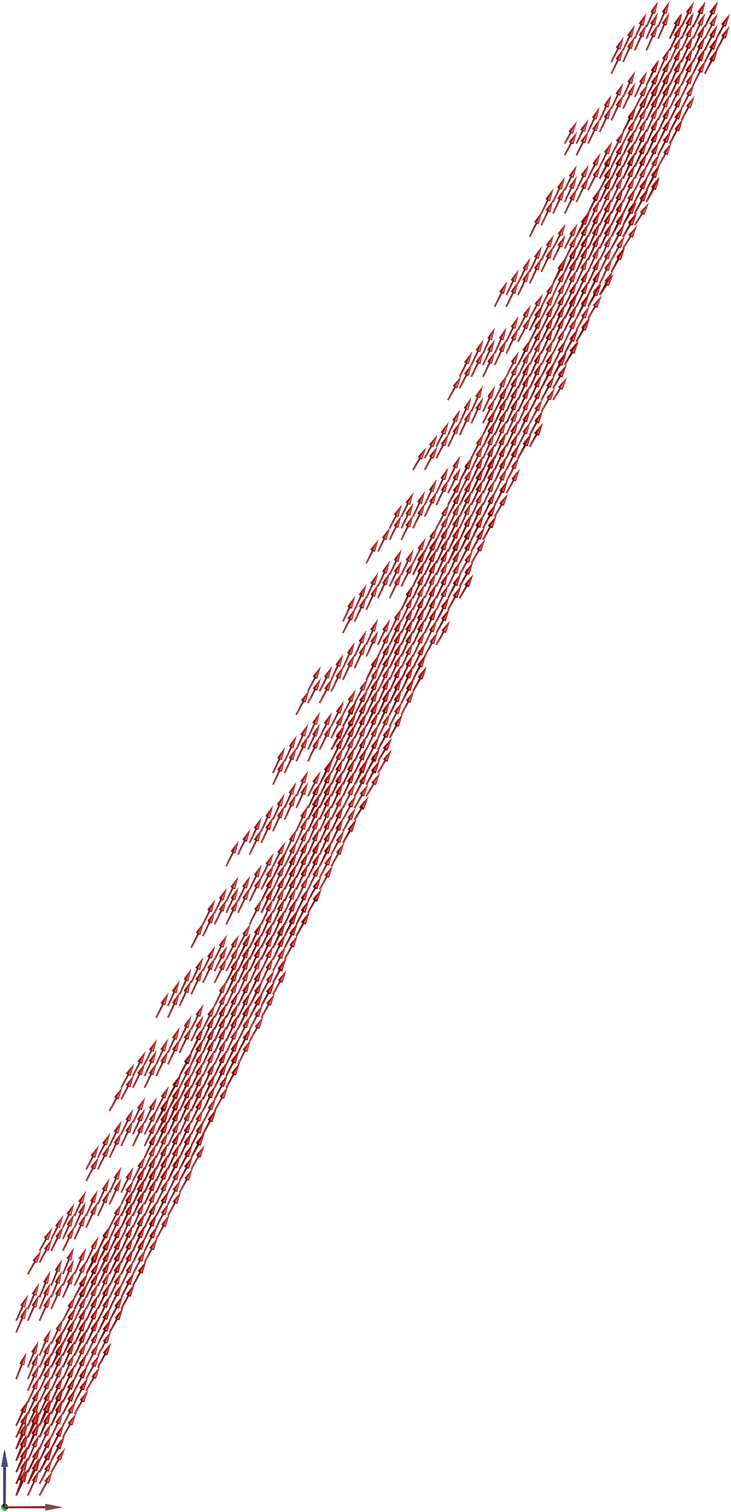

Figure S. 3. Parasagittal (XZ) plane projection of the 3D vector field from the synthetic fascicles based on the coordinate system of the aponeurosis. This visualization displays the fascicle orientations reconstructed using an isotropic 2 mm user-defined voxel size. Each red arrow represents the final 3D vectors derived from the hybrid tracking algorithm.

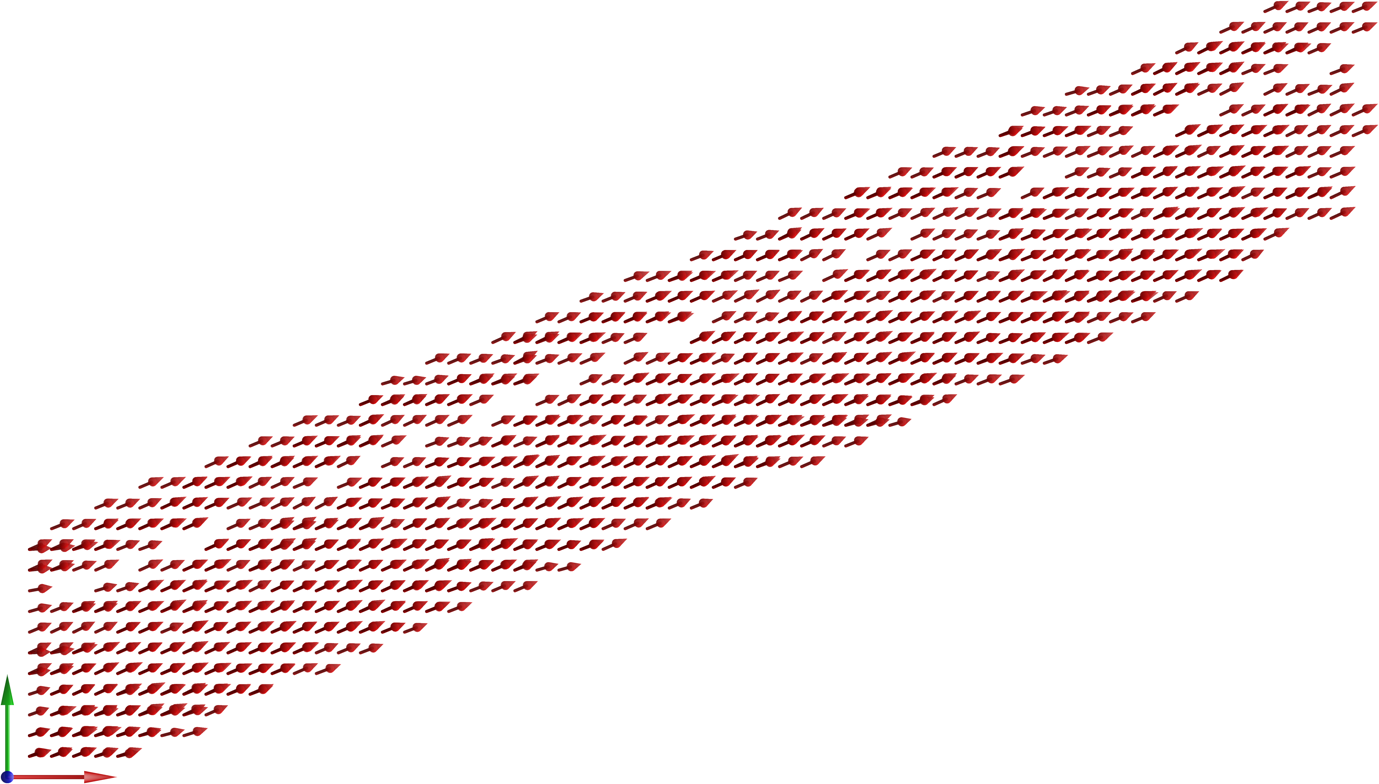

Figure S. 4. Frontal or medio-lateral (XY) plane projection of the 3D vector field from the synthetic fascicles based on the coordinate system of the aponeurosis. This visualization displays the fascicle orientations reconstructed using an isotropic 2 mm user-defined voxel size. Each red arrow represents the final 3D vectors derived from the hybrid tracking algorithm.

3DUS scan information

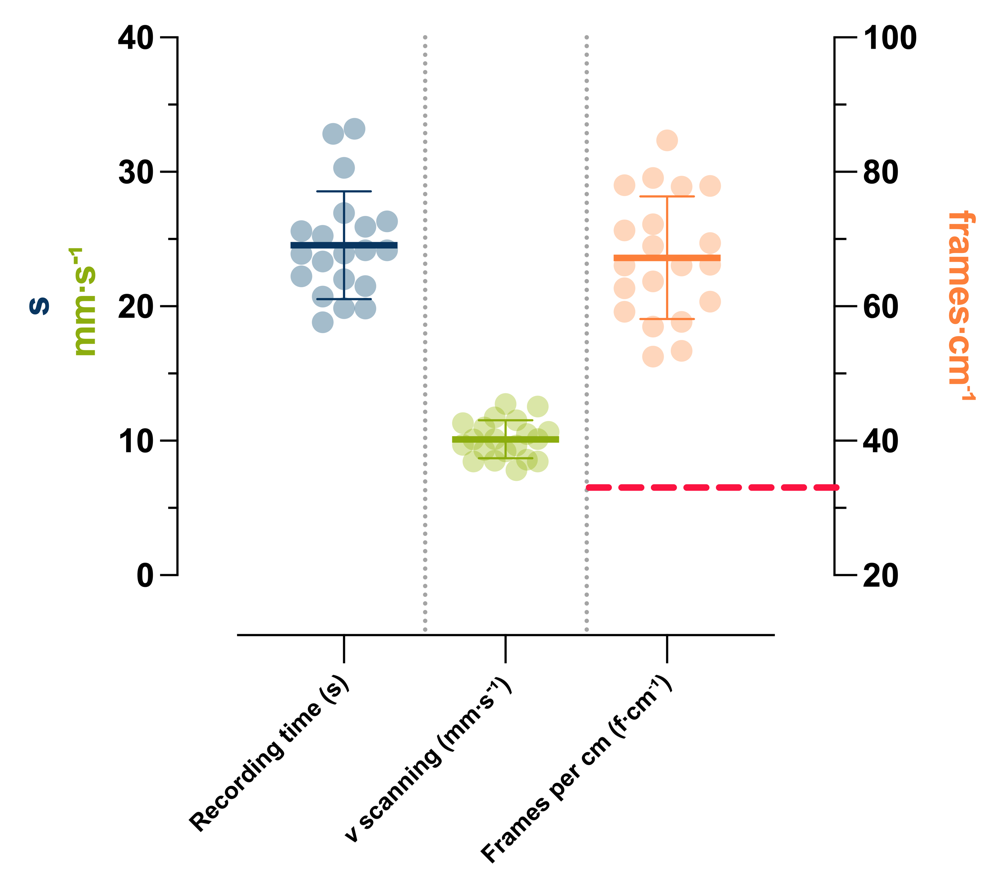

Figure S. 5. Summary of the recording time (s), velocity (v, mm/s) of the scan, and the frames obtained per cm (f/cm) during the 3DUS scans. The dots represent individual values, while the thick and thin horizontal bars indicate the mean ± standard deviation from all scans, respectively. The horizontal dashed red line in the right column denotes the “critical” threshold of 33 frames per cm defined by Nikolaev et al., 2022 required for proper voxel reconstruction.

Algorithm steps

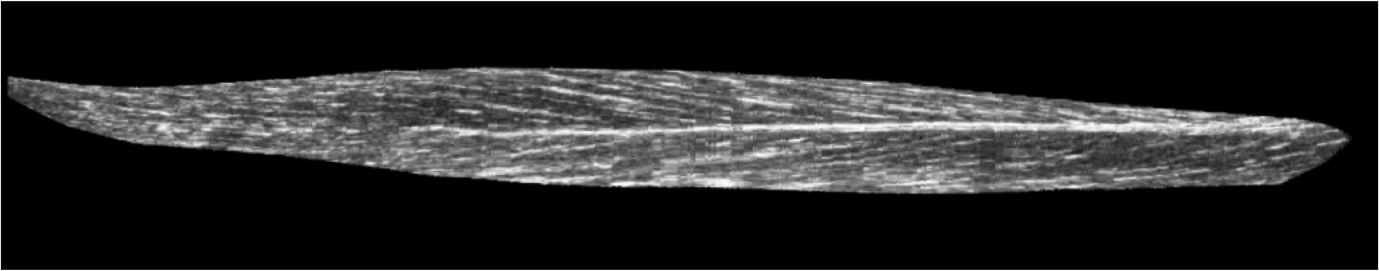

Figure S. 6. Example of a reconstructed parasagittal (XZ) plane slice showing fascicles within TA’s superficial and deep compartments (separated by the central aponeurosis) after affine transformation of the 3DUS data into the PCA-defined anatomical (aponeurosis-based) reference system.

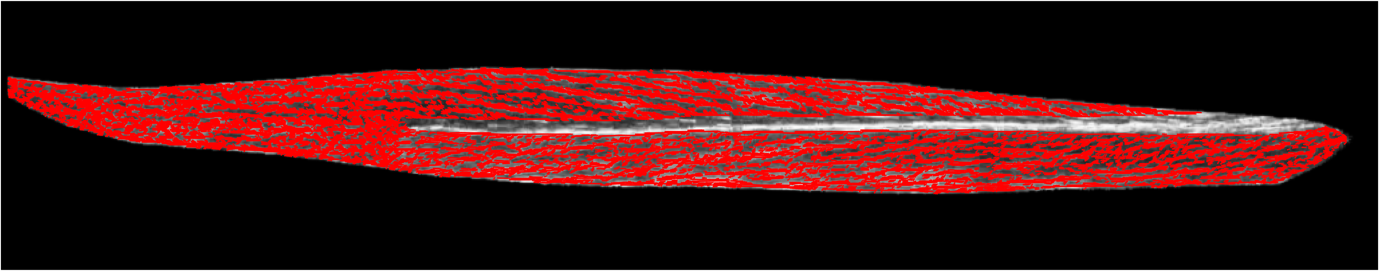

Figure S. 7. Example of the wavelet-based detection of fascicle orientations in the same parasagittal (XZ) slice as in Figure S. 6 using a 15x15 pixel size.

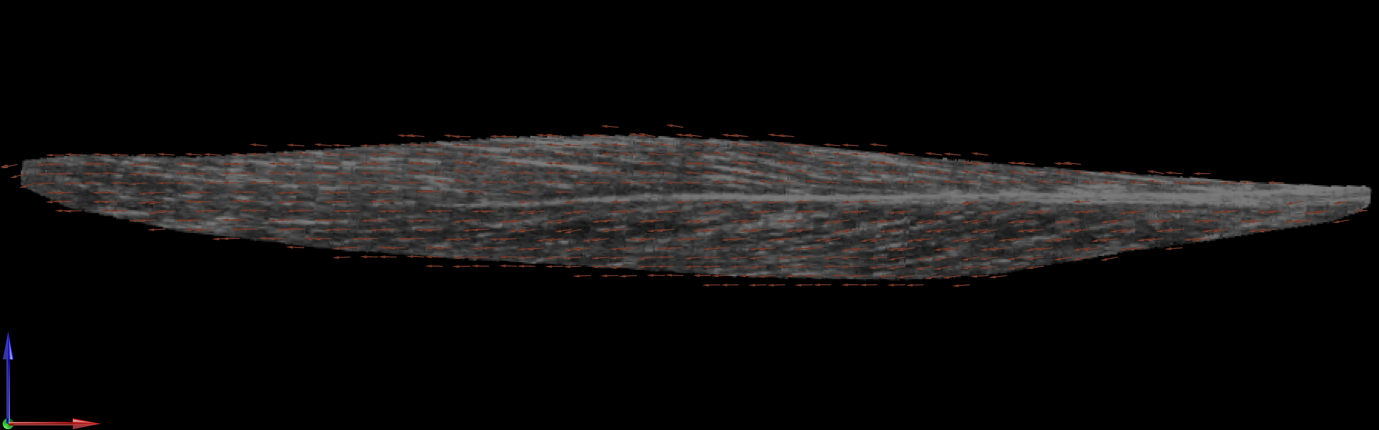

Figure S. 8. Example of the reconstructed 3D fascicle vector field projected in the same parasagittal (XZ) slice as in Figure S. 6 using an isotropic 3 mm user-defined voxel size.

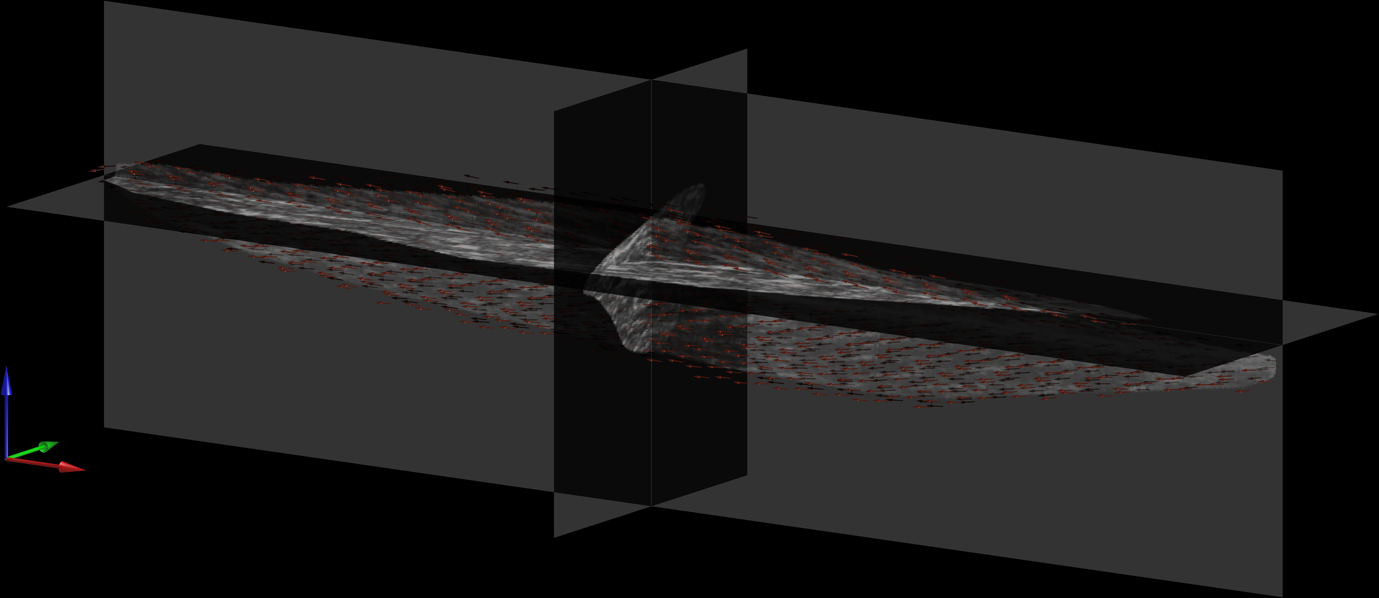

Figure S. 9. Reoriented parasagittal-plane dominant view of the reconstructed 3D fascicle vector field from Figure S. 8 based on ~20° rotation about the vertical axis to predominantly show the sagittal (XZ) plane projections of the fascicle vectors.

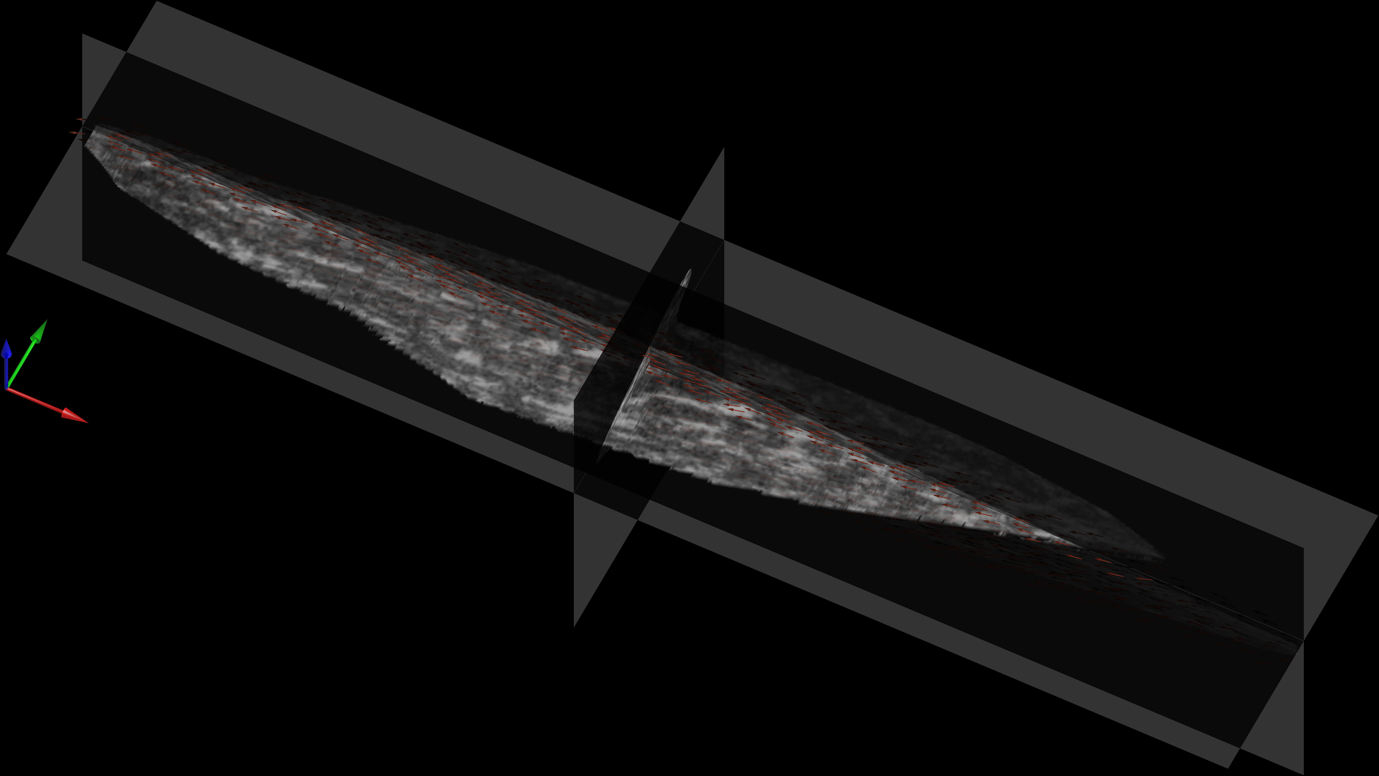

Figure S. 10. Reoriented frontal-plane dominant view of the reconstructed 3D fascicle vector field from Figure S. 8 based on a 20° rotation about the longitudinal axis to predominantly show the frontal (XY) plane projections of the fascicle vectors. This view highlights that local fascicle vectors align with the underlying “white line-like” structures.

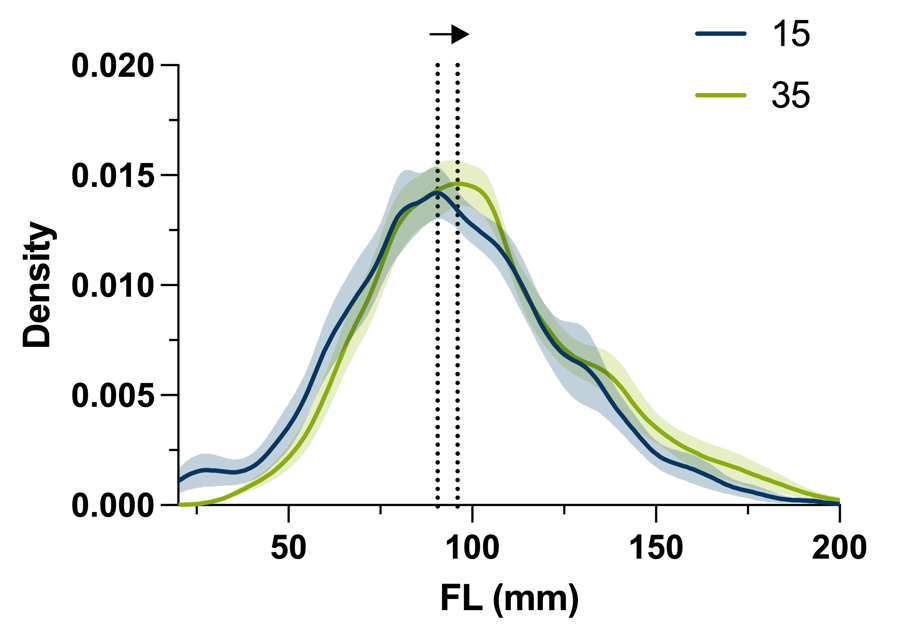

Figure S. 11. Probability density distributions of the tibialis anterior muscle fascicle lengths retrieved from 3DUS at two ankle angles. The solid lines represent the mean kernel density estimate at 15° (blue) and 35° (green) plantar flexion across all participants’ legs. The shaded regions indicate the standard error of the mean. Fascicle lengths were computed as the cumulative three-dimensional Euclidean distance along each reconstructed 3D vector until termination at the segmented central aponeurosis or muscle belly.

3D wire phantom data

To see how well our hybrid fascicle detection algorithm could handle more complex geometries than simple straight lines, we ran additional tests using 3D ultrasound data collected on three pairs of twisted wires submerged in a water bath. The three wires were soldered and twisted relative to a small rectangular base (a perforated plate originally used for soldering fine wires). The base served as a simulated aponeurosis and provided a fixed reference to objectively define the three main anatomical axes using a weighted principal component analysis, like with our current aponeurosis analysis approach with in vivo TA scans. The wires themselves curved and twisted throughout 3D space much more drastically than real muscle fascicles, making them an ideal way to stress our hybrid algorithm and see the limits of its performance. The hybrid approach followed the wire orientations well based on image reconstructions from multiple 2D planes, showing that the algorithm can accurately detect orientations of tubular structures that follow a complicated 3D path. However, once the local orientation reached around 20°, the orientation was underestimated by our hybrid approach, especially in the frontal (XY) plane (see Figure S17).

It is worth noting that the largest wire was not completely captured by the ultrasound image field of view during the 3DUS scan, but the data still serve as additional evidence of our hybrid algorithm’s robustness. Additionally, only scanning a few isolated wires further stressed our hybrid algorithm because of less local redundancy to help stabilize the orientation estimates, which is less of an issue compared with scanning muscle tissue, which has many parallel fascicles and thus increased local redundancy.

The reconstructed 3D vector fields from Figures S. 14-16 were generated using an isotropic user-defined voxel size of 3 mm. The reason 3 mm was used is to show the issue discussed within the main text; that is, non-physiological curvatures may be underestimated by our hybrid fascicle detection algorithm.

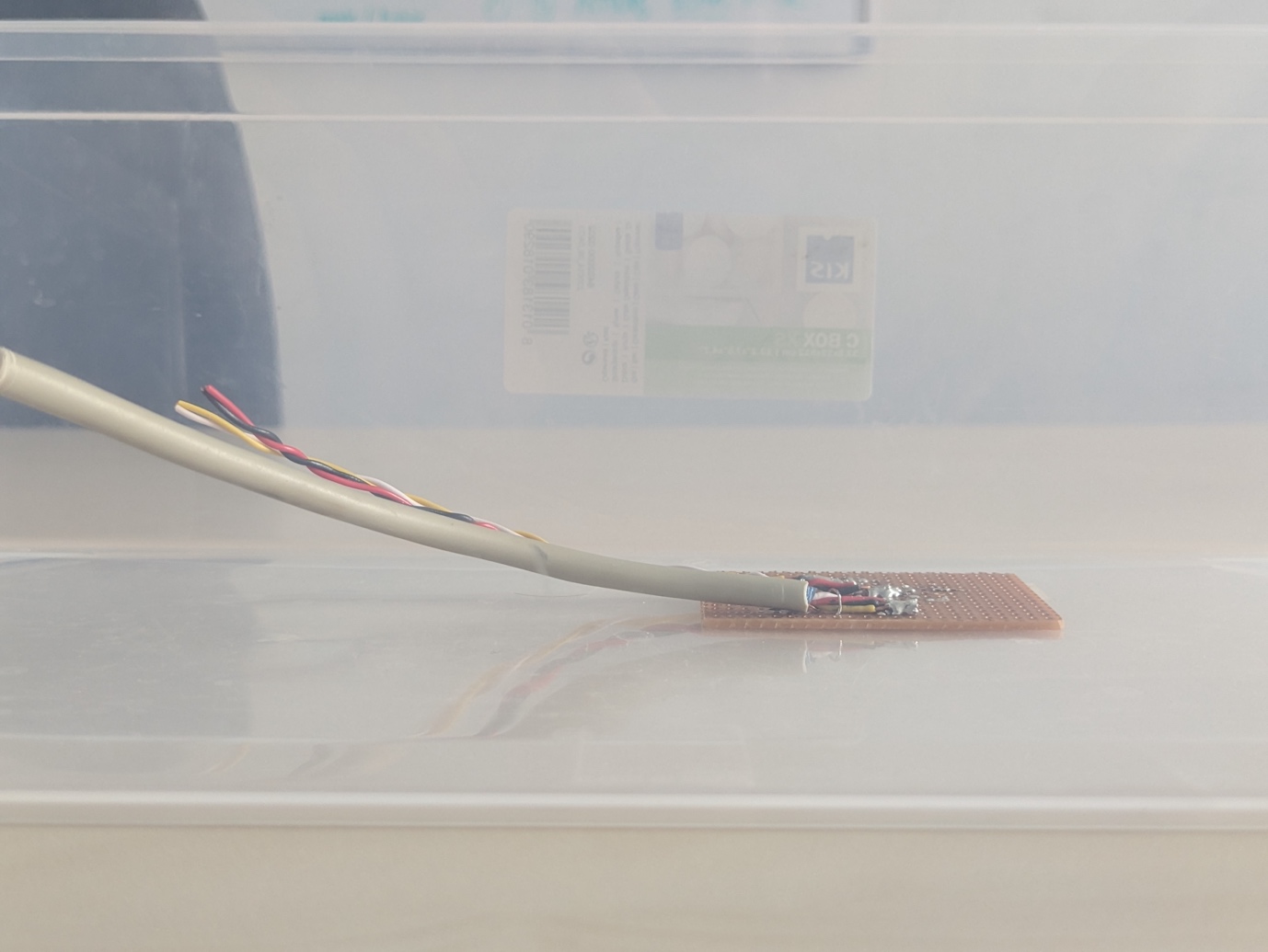

Figure S. 12. Photo of the phantom wires used from a sagittal plane perspective.

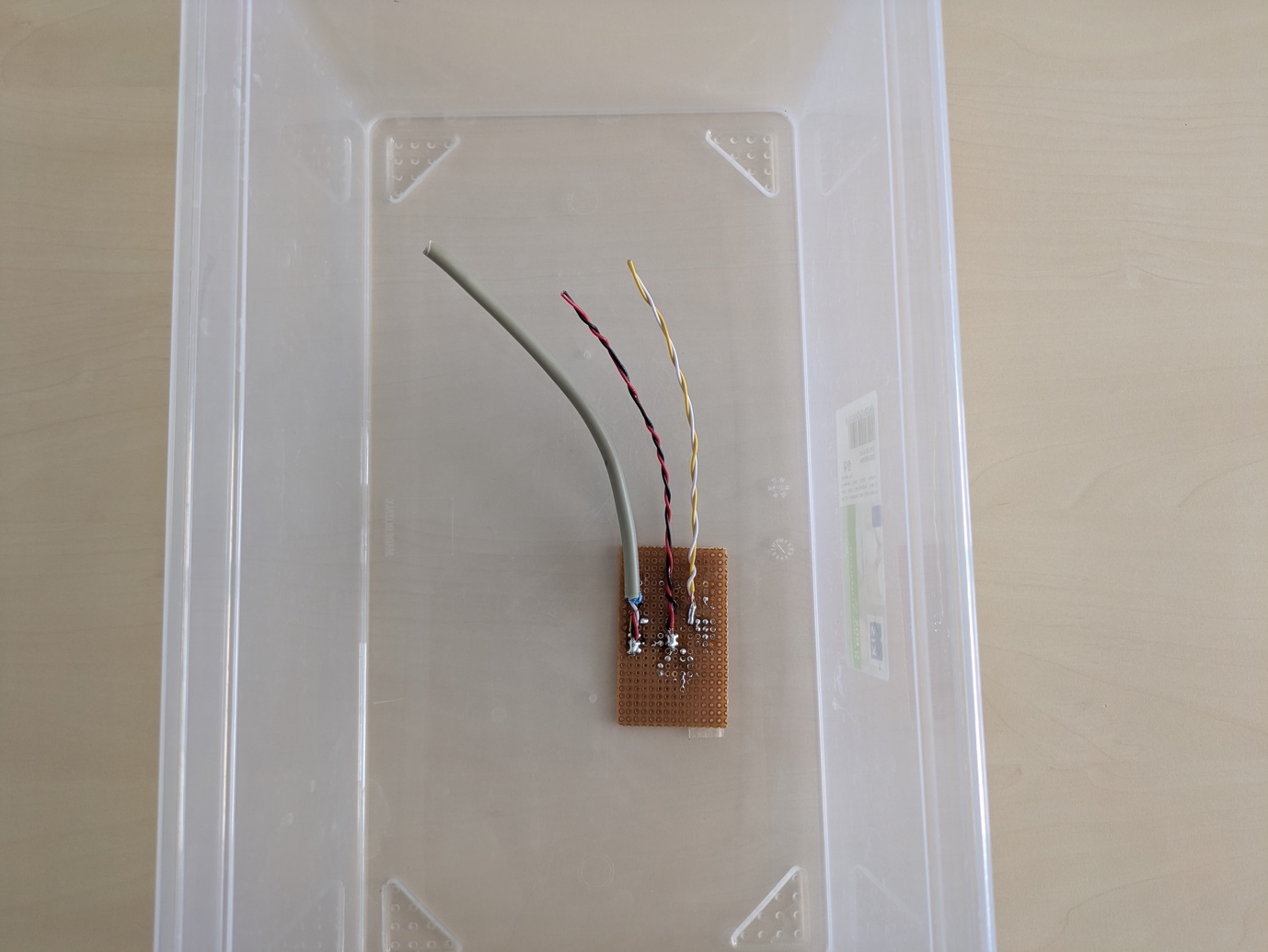

Figure S. 13. Photo of the phantom wires used from a frontal plane (birds-eye) perspective.

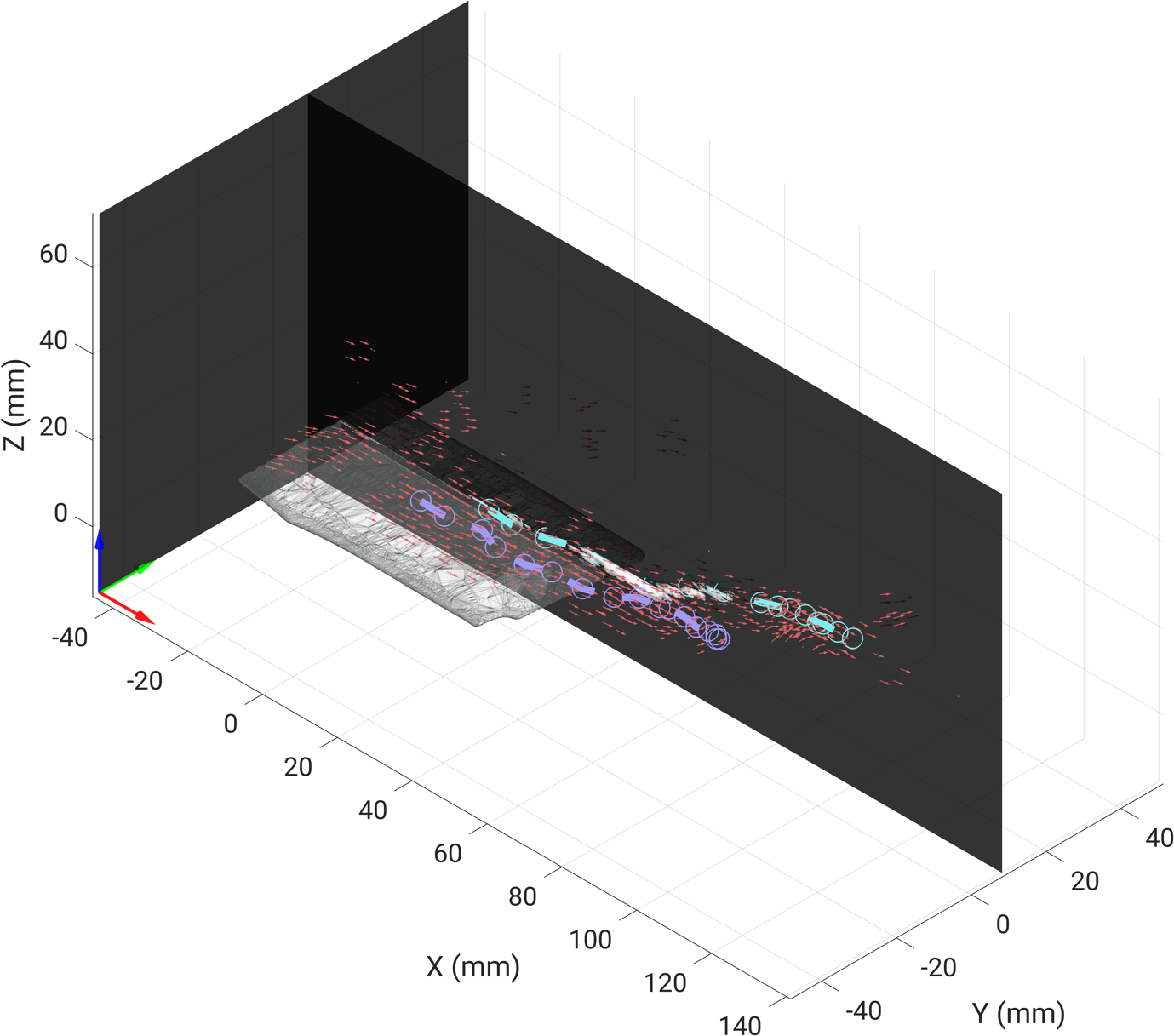

Figure S. 14. Combined frontal and sagittal plane view of the reconstructed 3D fascicle vector field from the phantom wire volume from 3DUS data. The light grey planar structure is the simulated aponeurosis, represented by the soldering base. The solid dashed light blue and violet lines show two fully manually digitized wires, while the red arrows represent the final 3D vectors derived from the hybrid tracking algorithm. For visual clarity, only a subset of 3D vectors is displayed.

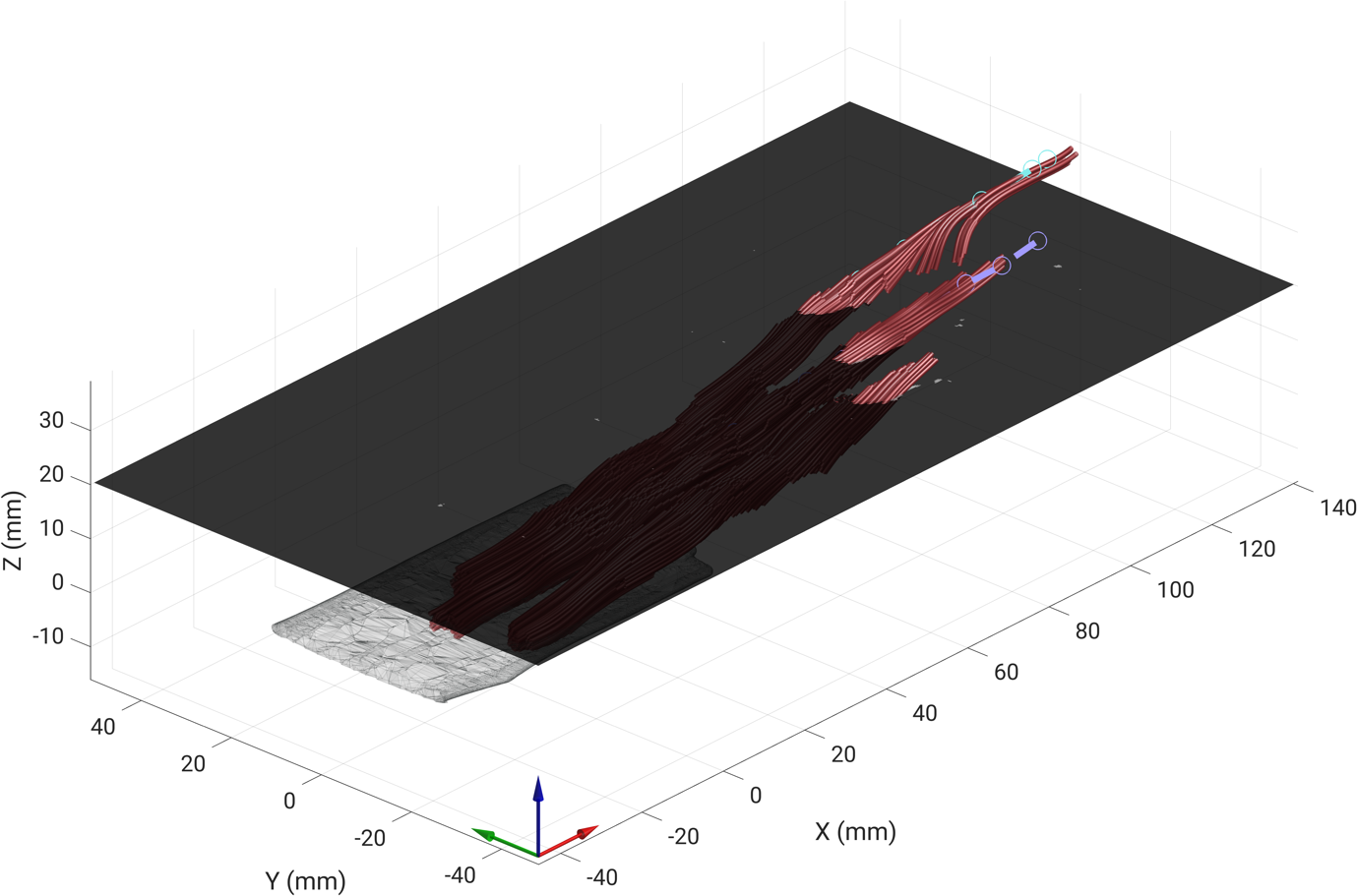

Figure S. 15. Combined frontal and sagittal plane view of the reconstructed 3D phantom wire paths. The XY slice at approximately 20 mm on the vertical axis (Z) marks the region where the algorithm begins to underestimate the frontal (XY) orientation component, resulting in a slight shortening of the 4^th^ order Runge-Kuttha tractography paths. This illustrates the observed limitation in orientation detection for local curvatures exceeding ~20°.

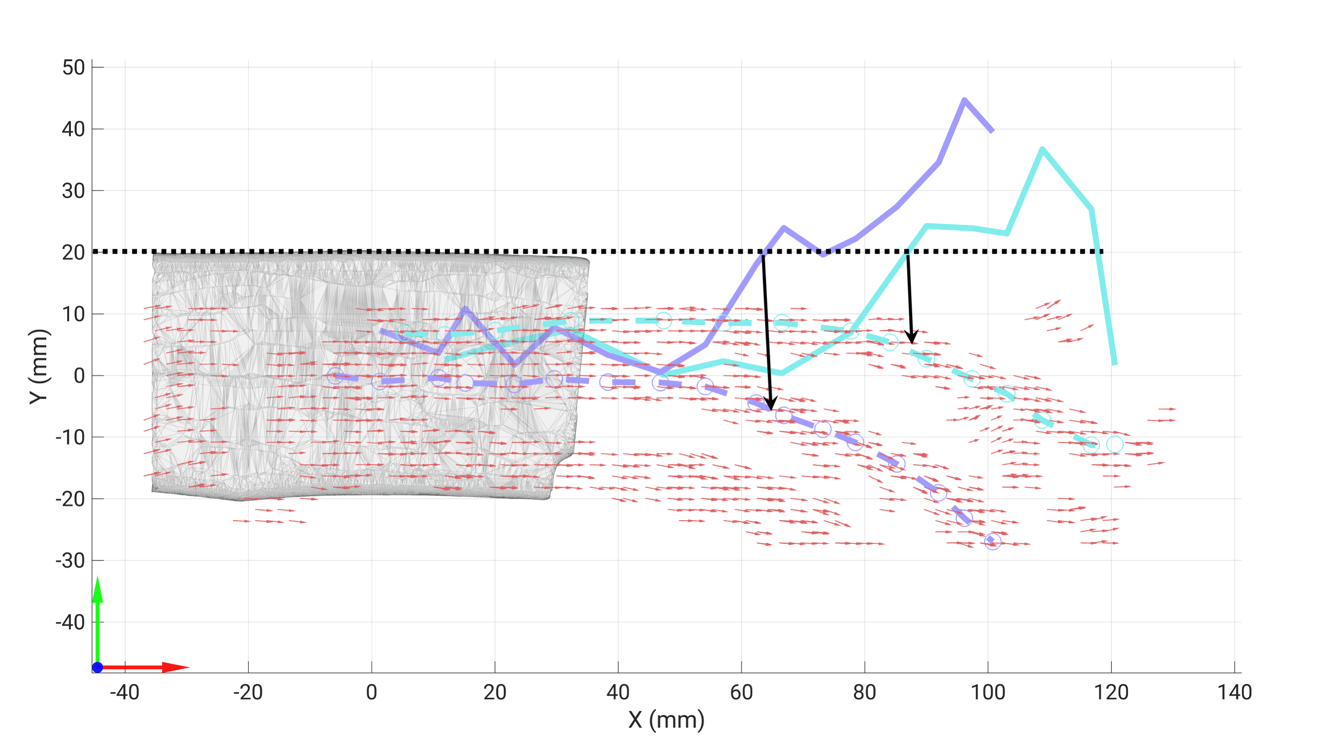

Figure S. 16. Frontal (XY) plane view of the reconstructed 3D vector field (red arrows). The two solid dashed lines represent the manually digitized local paths (location in mm) in the XY plane for the two fully-visible wires. The two solid lines represent the local orientations (in degrees) between the Y and X coordinates using the arctangent function (atan2d) applied to the manually digitized landmarks in Stradview. The black horizontal dashed line indicates the 20° orientation threshold, and the two black arrows highlight the detected vectors corresponding to the critical threshold where the frontal (XY) plane orientation begins to be underestimated.
